# A genomic and morphometric analysis of alpine bumblebees: ongoing reductions in tongue length but no clear genetic component

**DOI:** 10.1101/2021.03.17.435598

**Authors:** Matthew J. Christmas, Julia C. Jones, Anna Olsson, Ola Wallerman, Ignas Bunikis, Marcin Kierczak, Kaitlyn M. Whitley, Isabel Sullivan, Jennifer C. Geib, Nicole E. Miller-Struttmann, Matthew T. Webster

## Abstract

Over the last six decades, populations of the bumblebees *Bombus sylvicola* and *Bombus balteatus* in Colorado have experienced decreases in tongue length, a trait important for plant-pollinator mutualisms. It has been hypothesized that this observation reflects selection resulting from shifts in floral composition under climate change. Here we used morphometrics and population genomics to determine whether morphological change is ongoing, investigate the genetic basis of morphological variation, and analyze population structure in these populations. We generated a genome assembly of *B. balteatus*. We then analyzed whole-genome sequencing data and morphometric measurements of 580 samples of both species from seven high-altitude localities. Out of 281 samples originally identified as *B. sylvicola*, 67 formed a separate genetic cluster comprising a newly-discovered cryptic species (“incognitus”). However, an absence of genetic structure within species suggests that gene flow is common between mountains. We found a significant decrease in tongue length between bees collected between 2012-2014 and in 2017, indicating that morphological shifts are ongoing. We did not discover any genetic associations with tongue length, but a SNP related to production of a proteolytic digestive enzyme was implicated in body size variation. We identified evidence of covariance between kinship and both tongue length and body size, which is suggestive of a genetic component of these traits, although it is possible that shared environmental effects between colonies are responsible. Our results provide evidence for ongoing modification of a morphological trait important for pollination and indicate that this trait likely has a complex genetic and environmental basis.

## INTRODUCTION

The importance of pollinators for ecology and the maintenance of biodiversity is widely recognized (Goulson, Nicholls, Botías, & Rotheray, 2015). Bumblebees are large-bodied, mainly cold-adapted species with broad significance for agricultural production and plant-pollinator networks in the wild (Goulson, 2003). Many bumblebee species are experiencing range shifts or declines due to climate change or habitat loss (Cameron et al., 2011; Cameron & Sadd, 2020; Goulson, Lye, & Darvill, 2008a; Soroye, Newbold, & Kerr, 2020). In particular, a general worldwide trend for species to shift to higher elevations and latitudes has been documented, with losses occurring at a faster rate than expansions (Kerr et al., 2015; Marshall et al., 2020). Arctic and alpine species are particularly threatened by these trends, and can be considered “canaries in the coal mine” for detecting early effects (Elsen & Tingley, 2015).

In addition to range shifts there is evidence that the morphology of certain populations of bumblebees has shifted in recent decades. Miller-Struttmann et al. (2015) compared morphometric measurements of contemporary samples and historical specimens of the species *B. balteatus* and *B. sylvicola* from the same high-elevation alpine localities in Colorado. These species historically comprise the majority of bumblebees in these locations (Miller-Struttmann et al., 2015). They reported a significant decrease in tongue length in both species in a period between 1966 and 2014. The cause of this was inferred to be a general decrease in floral abundance resulting from warmer temperatures and drier soils resulting from climate change. These conditions could favor more generalist foraging strategies, which several studies indicate are performed more efficiently by bees with shorter tongues (Goulson & Darvill, 2004; Goulson, Hanley, Darvill, Ellis, & Knight, 2005; Goulson, Lye, & Darvill, 2008b; Heinrich, 2004; Huang, An, Wu, & Williams, 2015). Continued monitoring of these populations is important to understand the causes of these morphological shifts and their ecological consequences. It is unknown whether the trend towards shorter tongues has continued in more recent years.

One explanation for the observed morphological shifts in *B. balteatus* and *B. sylvicola* in Colorado is that natural selection for shorter tongues has occurred (Miller-Struttmann et al., 2015). However, it also possible that the observed changes are a plastic response to environmental change (Merilä & Hendry, 2014). It is unknown whether this trait has a genetic component that could be acted on by natural selection. Uncovering the size of the genetic contribution to this trait and the number and identity of any genetic loci involved would be an important step towards understanding the underlying cause of the morphological changes. The feasibility of genome sequencing on a population scale has enabled studies of quantitative genetics (Gienapp et al., 2017) and genome-wide association studies (Santure & Garant, 2018) in wild populations, allowing these goals to be pursued.

The genetic structure of populations is an important factor for interpretation of genotype-phenotype correlations and the cause of morphological shifts. We previously presented a genome assembly of *B. sylvicola* and analyzed genome-wide variation by resequencing 281 samples (Christmas et al., 2021). This identified the presence of a previously-undetected cryptic species, which we gave the preliminary name *Bombus incognitus*. We refer to this species hereafter simply as “incognitus” to reflect the fact that a formal taxonomic description is not currently available. This species was identified morphologically as *B. sylvicola* but forms a distinct genetic cluster. It is therefore possible that changes in tongue length in the combined populations of these species could be brought about by changes in the relative abundance of each species, although this has not previously been investigated. Additional population structure within species could lead to a similar effect, particularly if genetic structure associated with morphology exists. An understanding of genetic structure and spatial connectivity between populations in different locations is also needed to predict how they will evolve under climate change and to define the conservation value of subpopulations (Pauls, Nowak, Bálint, & Pfenninger, 2013; Razgour et al., 2019).

The degree of gene flow in bumblebee populations is determined by the distances that reproductive individuals (males and queens) travel in order to mate and establish new nests (Heinrich, 2004; Woodard et al., 2015). Genetic methods have been used to investigate population structure of several species of bumblebees across a variety of habitats in the UK, continental Europe and continental USA (Darvill, Ellis, Lye, & Goulson, 2006; Ellis, Knight, Darvill, & Goulson, 2006; Ghisbain et al., 2020; Jackson et al., 2018; Jha, 2015; Koch, Looney, Sheppard, & Strange, 2017; Woodard et al., 2015). A general finding is that populations of common species tend to exhibit very little structure in the absence of geographical barriers, even at continental scales. However, genetic differentiation can occur in rare or declining species, or when there are natural barriers (Woodard et al., 2015). Populations restricted to high elevations have been shown to exhibit higher genetic differentiation (Lozier, Strange, & Koch, 2013; Lozier, Strange, Stewart, & Cameron, 2011). However, the degree of fragmentation of populations of bumblebee species inhabiting high elevations in mountain ranges is not fully understood.

Genetic methods can also be applied to estimate nest density and foraging distances of bumblebees, by analyzing where genetically-related nestmates are caught. Such studies indicate that maximum foraging distances range from less than 100 m to over 10 km (Darvill, Knight, & Goulson, 2004; Geib, Strange, & Galen, 2015; Jha & Kremen, 2013; Knight et al., 2005; Woodard et al., 2015). These variables show large variation among species and are strongly dependent on landscape composition. Genome sequencing of multiple individuals has the potential to add greater resolution to our understanding of population density, structure and dispersal.

In this study we present a high-quality annotated genome assembly of the bumblebee species *B. balteatus* based on long-read Oxford Nanopore sequencing. We also conducted whole-genome sequencing of 299 samples of this species sampled from seven high-altitude localities spread across 150 km in Colorado. These localities have an average separation of 54 km between each other, and consist of alpine tundra separated by forest. We analyze these data together with the previously-published population genomic dataset of 284 samples of *B. sylvicola* and “incognitus” from the same localities (Christmas et al., 2021). We use these data to assess whole-genome variation and its connection to geographical and morphological variation in these populations, which were previously studied by Miller-Struttmann et al. (2015). We then analyzed morphological variation in tongue length and body size in context of previous studies (Miller-Struttmann et al., 2015) to determine whether the trend towards decreasing tongue length is ongoing. We also performed a genome-wide analysis of the genetic basis of morphological variation using both genome-wide association studies (GWAS) and genome-wide complex trait analysis (GCTA) (Yang, Lee, Goddard, & Visscher, 2011). The results are informative regarding the degree of connectivity among bumblebee populations, the genetic basis of morphological variation, and the mechanisms of morphological evolution.

Our study species have recently been subject to taxonomic revision. The species we refer to here as *B. balteatus* was previously described as occurring in both North America and Eurasia, but a recent study classified these geographically separated populations as two different species: *B. balteatus* in Eurasia and *Bombus kirbiellus* in North America (Williams et al., 2019). The species we refer to here as *B. sylvicola* was previously split into a North American form (*B. sylvicola*) and a Eurasian form (*Bombus lapponicus*). However, a recent analysis synonymized these two species as *B. lapponicus* (Martinet et al., 2019). We continue to use the previous names here to avoid confusion with comparison to previous ecological and evolutionary genetic studies.

## MATERIALS AND METHODS

### Sample collection

We collected samples identified as *B. balteatus* and *B. sylvicola* (Koch, Strange, & Williams, 2014) during the summer of 2017. Bumblebees were collected on seven different mountains in Colorado, USA, with an average distance of 54 km between mountains (greatest distance: 134 km, Quail Mountain – Niwot Ridge; shortest distance: 10km, Mount Democrat – Pennsylvania Mountain). The existence of “incognitus” was unknown to us when the bees were sampled but a large fraction of bees identified as *B. sylvicola* were subsequently assigned to “incognitus” on the basis of genetic analysis (Christmas et al., 2021). Three of these mountains (Mount Evans, Niwot Ridge, and Pennsylvania Mountain) were also sampled previously by Miller-Struttmann et al. (2015). Four additional mountains were sampled here (Boreas Mountain, Mount Democrat, Horseshoe Mountain and Quail Mountain) but were not sampled in the previous study. Each of the mountain locations comprised several neighboring sampling sites separated by short distances (mean = 1.1 km) and elevations (range: 3,473 – 4,012 m.a.s.l.) (full details in supplementary tables S1, S2).

We collected samples of foraging worker bees from each site using sweeping hand nets. We kept samples in Falcon tubes in cool boxes for transport. Species assignment was performed by observing standard morphological characters (Koch et al., 2014). We dissected each bee after placing it at −20°C for approximately 10 minutes.

### Morphological measurements

We measured intertegular distance and tongue length on all of the samples of *B. sylvicola*, “incognitus” and *B. balteatus* using scaled photographs of individual bee heads processed by ImageJ (Kearns & Inouye, 1993; Schneider, Rasband, & Eliceiri, 2012). Following netting, bees were anesthetized by chilling them on ice or in a freezer (depending on field conditions). Once anesthetized, the bees were removed from the vial, gently pressed prone to the table with the thorax parallel to the camera lens using flat forceps, and photographed with a metric ruler for scaling purposes. The distance between the tegulae was measured as a straight line between the edge of one tegula to the other tegula, running as near to perpendicular to the tegula as possible. The head was then immediately removed from the body and placed into an individual labeled vial and kept frozen until further characterization in the lab. Following the protocol used in Miller-Struttmann *et al*. (2015), the prementum and glossa were dissected from the head while the tissue was flexible, straightened and aligned as possible without damaging the tissue, and mounted on acid-free paper. The samples were then photographed with a ruler in the field of view under a microscope, and the traits measured (i.e., prementum length and glossa length) via ImageJ (Schneider et al., 2012). After photographing, we stored thoraces in 95% ethanol prior to DNA extraction. We visually characterized the diagnostic traits visible on each head, such as the shape of the malar space, the pile color between and antenna and above the ocelli, the size, color and location of the ocelli relative to the supraorbital line, and the presence of black pile on the scutellum. None of the morphological characters we measured could be used to distinguish between “incognitus” and *B. sylvicola*.

### Genome sequencing and assembly

We used the same sequencing strategy to produce high-quality genome assemblies for both *B. sylvicola* and *B. balteatus*. The *B. sylvicola* assembly was presented previously (Christmas et al., 2021). This involved combining data from Oxford Nanopore (ONT) and 10x Genomics Chromium technologies. For both species, we used DNA extracted from male bees sampled on Niwot Ridge and stored in 95% ethanol prior to DNA extraction. DNA from a single bee was used for each assembly. We performed DNA extraction using a salt-isopropanol extraction followed by size selection using magnetic bead purification to remove fragments < 1 kbp. ONT sequencing was performed using the MinION instrument. For *B. sylvicola*, we used two R9.4 flowcells and the RAD004 kit starting with 3-400 ng DNA per run. This resulted in a total of 9.4 Gbp of sequence data, in 2.5 million reads with a mean length of 3.7 kbp (Christmas et al., 2021). For *B. balteatus*, we used 1μg DNA in a modified LSK-108 one-pot protocol. 1.75 μl NEB FFPE buffer, 1.75 μl NEB Ultra-II End Prep buffer and 1 μl FFPE repair mix was added to 24 μl DNA and incubated at 20°C for 30 minutes. 2 μl End Prep enzyme mix was then added for blunting, phosphorylation and A-tailing (20 min. 20°C, 10 min. 65°C, 10 min. 70°C). AMX (20 μl), ligation enhancer (1 μl) and Ultra-II ligase (40 μl) was added for ligation at room temperature for 30 minutes. Clean-up and flowcell loading was done following the standard ONT LSK-108 protocol and the library was sequenced on one R9.4 flowcell. This resulted in 23.3 Gpb of sequence data, in 3.9 million reads with a mean length of 5.9 kbp.

We assembled the ONT reads using the following process: We first used downpore (Teutenberg, 2018/2020) with default parameters for adaptor trimming and splitting chimeric reads. We then assembled the trimmed reads using wtdbg2 (Ruan & Li, 2020) on default settings. We then ran two rounds of the consensus module Racon (Vaser, Sović, Nagarajan, & Šikić, 2017) and contig improvements using medaka v.0.4 (*Medaka*, 2017/2021), removing contigs of length < 20 kbp. We used Illumina short-read data from the population sequencing to perform two rounds of Pilon (Walker et al., 2014) to improve sequence accuracy, particularly around indels.

We also sequenced the same sample of each species using 10x Genomics Chromium. A 10x GEM library was constructed according to the manufacturer’s protocols. We normalized each library using qPCR and sequenced them together on a single lane of HiSeq 2500 using the HiSeq Rapid SBS sequencing kit version 2, resulting in 150 bp paired-end sequences. The linked reads were mapped to the assembly using Longranger v.2.1.4 (*10XGenomics/Longranger*, 2018/2021). We then ran Tigmint v1.1.2 (Jackman et al., 2018) to correct errors in the assembly detected by the read mappings. We identified contigs that contained mitochondrial genes, and were therefore likely fragments of the mitochondrial genome, by running a BLAST search *of B. impatiens* mitochondrial genes across the assembly using BLAST+ v2.9.0 (Camacho et al., 2009). Any contigs containing two or more mitochondrial genes located within the expected distance of each other based on their locations on the mitochondrial genome were removed from the assembly to ensure the final assembly did not contain partially assembled mitochondrial genome sequence. We also removed all contigs shorter than 10 kbp.

### Genome analysis

BUSCO v3.0.2b (Simão, Waterhouse, Ioannidis, Kriventseva, & Zdobnov, 2015) was used to assess the assembly completeness using the hymenoptera_odb9 lineage set and species *B. impatiens*. We ran RepeatMasker v.4.1.0 (Smit, Hubley, & Green, 2015) on each genome assembly to characterize genome repeat content, including interspersed repeats and low complexity sequences. Chromosome-level assemblies of species from five diverse bumblebee subgenera indicate that non-parasitic clades have a stable 18 chromosome karyotype (Sun et al., 2020). We therefore performed whole-genome synteny alignments between the *B. terrestris* chromosome-level genome assembly (Sadd et al., 2015) (downloaded from NCBI; BioProject PRJNA45869) and our assemblies using Satsuma v.3 (Grabherr et al., 2010) to arrange the contigs from our assemblies onto 18 pseudochromosomes.

### RNAseq and Annotation

We used the Nextflow pipelines available at https://github.com/NBISweden/pipelines-nextflow for genome annotation. The pipelines, in turn, depend on two additional annotation-specific toolkits, AGAT and GAAS, available at https://github.com/NBISweden/AGAT and https://github.com/NBISweden/GAAS respectively. Both repositories contain extensive documentation and installation instructions. As reference proteins database, we used the SwissProt/UniProt (561,356 curated proteins, downloaded 2019-11). The parameter files stated below are available at https://github.com/matt-webster-lab/alpine_bumblebees.

We performed RNAseq on a single lane of Illumina HiSeq 2500 using samples from four body parts from a single sample of *B. balteatus* (abdomen, head, legs and thorax), which generated 42 Gbp of reads. We performed quality control of the data using FastQC (Andrews, 2017/2021) and read trimming using trimmomatic (v. 0.39, trimmomatic.params). We next generated a guided assembly with Hisat2 (Kim, Paggi, Park, Bennett, & Salzberg, 2019) and Stringtie (Pertea et al., 2015) using the AnnotationPreprocessing.nf (parameters as in AnnotationPreprocessing_params.conf) TranscriptAssembly.nf Nextflow pipeline with parameters as in the TranscriptAssembly_params.config file. In addition to the guided assembly, we performed a de novo assembly for reads from each of the four tissues using Trinity v. 2.0.4 (Grabherr et al., 2011) (--seqType fq --SS_lib_type RF).

Next, we constructed a species-specific repeat library using RepeatModeler package (v. open-1.0.8; -engine ncbi -pa 35). Since the de novo identified repeats may still include parts of protein-coding genes, we perform an additional filtering step and remove from the database the repeats that can be found in a comprehensive set of known proteins. We remove all repeats that match to a pre-computed set non-transposable proteins from our protein reference database. We used custom script (filter_repeats.bash) for filtering. To identify the location of repeats from this library in the genomes, we used RepeatMasker (open-4.0.9).

We used the MAKER (Cantarel et al., 2008) package to compute gene builds. An evidence-guided gene build was computed by MAKER (-fix-nucleotides) constructing gene models from these reference proteins and from the transcript sequences aligned to the genome assembly (parameters in evidence_build.maker_opts.ctl). We also prepared an *ab initio* build using the AbInitioTraining.nf pipeline and AbInitioTraining_params.config as the parameters file. Both the evidence-guided and the ab initio builds were than merged using MAKER (hybrid_build_maker_opts.ctl). Finally, we performed functional annotation using the FunctionalAnnotation.nf pipeline with FunctionalAnnotation_params.config as the parameter file.

### Population sequencing, read mapping and variant calling

We extracted DNA from the thoraces of 299 worker bees of *B. balteatus* collected from across the seven sampling sites using the Qiagen Blood and Tissue kit. We prepared dual-indexed libraries using the Nextera Flex kit and performed sequencing on an Illumina HiSeq X to produce 2 x 150 bp reads, using an average of 36 samples per lane. We mapped reads to the *B. balteatus* reference using the mem algorithm in BWA (Li & Durbin, 2009). We performed sorting and indexing of the resultant bam files using samtools (Li et al., 2009) and marked duplicate reads using Picard. We used the Genome Analysis Toolkit (GATK) to call variants (McKenna et al., 2010). We first ran HaplotypeCaller using default parameters on the bam file of each sample to generate a gVCF file for each sample. We then used GenomicsDBImport and GenotypeGVCFs with default parameters to call variants for all *B. balteatus* samples together. We applied a set of hard filters using the VariantFiltration tool to filter for reliable SNPs using the following thresholds: QD < 2, FS > 60, MQ < 40, MQRankSum < −12.5, ReadPosRankSum < −8 (see the GATK documentation for full descriptions of each filter). Only biallelic SNPs were considered for downstream analysis. Samples from *B. sylvicola* and “incognitus” samples were analyzed in the same way and are presented in Christmas *et al*. (2021).

### Relatedness and population structure

We estimated the pairwise kinship coefficient between all samples within each species using the method described in (Manichaikul et al., 2010) implemented in the --relatedness2 option of vcftools v0.1.16 (Danecek et al., 2011). Worker bees from the same bumblebee colony are expected to have a relatedness of 0.75 under the assumption of monandry, which is reasonable for the study species (Estoup, Scholl, Pouvreau, & Solignac, 1995). The kinship coefficient (φ) predicted from this relationship is 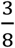 (the probability that two alleles sampled at random from two individuals are identical by descent). We estimated the lower cutoff for the inference criteria to infer this level of relatedness using powers of two as described in (Manichaikul et al., 2010), providing an estimate of 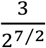.

We tested for correlations between relatedness and geographic distance for each species using Mantel tests with the mantel function in the ‘ecodist’ package in R v4.0.2. Coordinates for each sample were converted to geographic distances using the R package ‘geodist’ and these were compared to the matrix of kinship coefficients described above. We used the mantel.correlog function to produce mantel correlograms and assess whether correlations were significant over different geographic distances. We performed these analyses after excluding sisters from the datasets to ensure any patterns we observe were not driven by an effect of sampling individuals from the same colony.

We thinned the VCF files to 1 SNP every 10 kbp using the --max-missing 1 and --thin flags in vcftools v.0.1.16 to ensure no missing data and to assess population structure and relatedness more efficiently. We calculated p-distance matrices for *B. balteatus* samples and *B. sylvicola* and “incognitus” samples combined from the thinned vcf files using VCF2Dis (https://github.com/BGI-shenzhen/VCF2Dis) and then constructed neighbor-joining trees using the ‘ape’ package in R v4.0.2. We converted the thinned vcf files to plink format and performed principal components analyses using the ‘adegenet’ package in R v4.0.2. We performed PCAs on all samples as well after removing one of each pair of sisters identified via the kinship analysis to ensure that relatedness did not heavily influence any clustering. We used admixture v1.3.0 (Alexander, Novembre, & Lange, 2009) to assess the most likely number of genetic clusters within each species. We ran admixture for K=1 through to 10, with 10 iterations per K value and used the calculated cross-validation (cv) errors to assess the most suitable values of K, with low cv-error indicating higher support.

### Statistical morphometric analysis

We combined the phenotypic measurements made in this study with those from previous collections published in Miller-Struttmann et al. (2015). These measurements are intertegular distance, glossa length, prementum length, and total tongue length (equal to glossa length plus prementum length). This gave us three time periods to compare amongst: ‘2017’ (samples collected for this study), ‘2012-2014’, and ‘1966-1980’ (both sets published in Miller-Struttmann et al., 2015). We performed statistical analysis of variation in glossa length, prementum length, and intertegular distance in R v4.0.2. We investigated correlations between all traits for each species in the 2017 set using the ‘cor.test’ function, which calculates a Pearson’s product moment correlation coefficient.

Within our 2017 dataset, we compared glossa lengths between the three species by fitting linear models of species against glossa length with intertegular distance included as a random effect to account for body size using the lm function. We then calculated least-squares mean glossa lengths from this model using the lsmeans function and tested for significant differences using the Tukey method for p-value adjustment for multiple testing. We employed a similar approach for comparing glossa lengths between our dataset and historical datasets for each species. Here we used a linear mixed effect model framework with the ‘lmer’ function in the lme4 package in R v4.0.2, where mountain and intertegular distance were included as random effects in each model. Significance of each model was assessed using analysis of variance with the ‘anova’ function. We calculated least-squares mean glossa lengths from each model using the lsmeans function and tested for significant differences using the Tukey method for p-value adjustment for multiple testing.

We analyzed variation in intertegular distance between species and geographic locations for our 2017 dataset using analysis of variance (‘aov’ function) and assessed significance of comparisons using Tukey’s honestly significant difference (HSD) post hoc test (‘TukeyHSD’ function). For comparisons of intertegular distance between years, we fitted linear mixed effect models and included mountain as a random effect, then compared least-squares means from the models using the Tukey method for p-value adjustment for multiple testing.

### Genome-wide association study

We carried out mixed linear model-based association analyses, implemented in GCTA-MLMA (Yang et al., 2011; Yang, Zaitlen, Goddard, Visscher, & Price, 2014), to identify correlations between allele frequencies and trait variation across the genome for *B. balteatus* and *B. sylvicola* samples separately (“incognitus” samples were left out of this analysis due to too few samples). First, we created a genetic relationship matrix (GRM) from the genome-wide SNPs using the --make-grm function in GCTA. We removed one sample from each pair of sisters by setting a relatedness cut-off of 0.67 using the ‘--grm-cutoff’ flag. We ran the analyses for three traits per species: intertegular distance (ID), glossa length (GL), and the residuals from a linear regression of GL~ID to control for body size. The GRM was included as a polygenic/random effect. The analysis assumes phenotypes are normally distributed and so samples were removed in the tails of the distributions to meet this assumption for all traits except intertegular distance in *B. sylvicola*, which was normally distributed. For *B. balteatus*, this resulted in 279, 291, and 262 samples for the glossa length, intertegular distance, and residuals analyses respectively. For *B. sylvicola* there were 176, 192, and 152 samples for the glossa length, intertegular distance, and residuals analyses respectively. We assessed divergence of p-values from neutral expectations using QQ-plots and carried out Bonferroni correction of p-values using the ‘p.adjust’ function in R v4.0.2 to assess genome-wide significance.

### Genome-wide complex trait analysis

We used GCTA (Yang et al., 2011) to perform genome-wide complex trait analysis. We estimated the phenotypic variance explained by the genome-wide SNPs using genome-based restricted maximum likelihood (GREML). The analysis was performed separately for each species on measures of intertegular distance, glossa length, and glossa length with intertegular distance included as a covariate to control for body size. We ran each analysis twice, once using a GRM based on all samples and a second time with the same GRM produced above where one from each pair of sisters was removed. We calculated 95% confidence intervals around each estimate by multiplying the standard error by 1.96. We used R functions provided in (Wang & Xu, 2019) as well as the GCTA-GREML power calculator (Visscher et al., 2014) to perform power analysis and estimate the effect sizes we can expect to detect with our population datasets.

## RESULTS

### Generation of a highly-contiguous genome assembly of *B. balteatus*

We generated a *de novo* genome assembly for the species *B. balteatus* using Oxford Nanopore (ONT) long-read and 10x Chromium linked-read sequencing and compared this to the assembly for *B. sylvicola* that we recently published (Christmas et al., 2021). Assembly statistics for BBAL_1.0 (*B. balteatus*) and BSYL_1.0 (*B. sylvicola*) are presented in table 1, which includes statistics for the assemblies of *B. terrestris* (BTER_1.0) and *B. impatiens* (BIMP_2.2) for comparison. Total assembly sizes of 250.1 and 252.1 Mbp for *B. balteatus* and *B. sylvicola* respectively are both above the average assembly size of *Bombus* genomes assembled to date of 247.9 Mbp (22 genomes, range 229.8 – 282.1 Mbp) (Heraghty et al., 2020; Sadd et al., 2015; Sun et al., 2020). Contig N50s of 8.60 and 3.02 for BBAL_1.0 and BSYL_1.0 respectively are the longest and third longest contig N50s of all published *Bombus* assemblies, demonstrating these to be highly contiguous assemblies. Megabase-scale contig N50 are typical of long-read assemblies, such as the ones presented here and in Heraghty et al. (2020), and are significantly more contiguous than those based on short-read sequencing technologies such as BTER_1.0 and BIMP_2.2.

**Table 1:** Assembly metrics for *B. balteatus*, with three other published *Bombus* genomes included for comparison

| Species | *B. balteatus* | *B. sylvicola* | *B. terrestris* | *B. impatiens* |
| --- | --- | --- | --- | --- |
| Assembly | BBAL_1.0 | BSYL_1.0 | Bter_1.0 | BIMP_2.2 |
| Size (Mbp)<sup>†</sup> | 250.07 | 252.08 | 248.65<br>(236.38) | 246.86<br>(241.98) |
| Scaffolds (n) | - | - | 5,678 | 5,460 |
| Contigs (n) | 336 | 592 | 10,672 | 16,060 |
| Contig N50 (Mbp) | 8.60 | 3.02 | 0.08 | 0.06 |
| Contig L50 (n) | 12 | 28 | 890 | 54 |
| GC (%) | 37.64 | 38.23 | 37.51 | 37.76 |
| Repeat content (%) | 17.1 | 17.9 | 14.8 | 17.9 |
| Complete hymenoptera BUSCO genes (%)<sup>*</sup> | 99.0 | 98.2 | 96.9 | 98.3 |

<sup>\*</sup>BUSCO analysis using the OrthoDB v. 10, Hymenoptera dataset.
<sup>†</sup> Numbers in brackets represent total ungapped length

High genome completeness and accuracy is also reflected in the BUSCO analysis, with scores of 99.0% (98.7% single copy, 0.3% duplicated, 0.5% fragmented, 0.5% missing) and 98.2% (97.9% single copy, 0.3% duplicated, 0.4% fragmented, 1.4% missing) reported for BBAL_1.0 and BSYL_1.0 respectively. Whole genome alignments to the *B. terrestris* assembly (Sadd et al., 2015) resulted in 93.9% and 91.1% of the *B. balteatus* and *B. sylvicola* genomes being placed on 18 pseudochromosomes. Our annotation pipeline annotated 11,711 and 11,585 genes in BBAL_1.0 and BSYL_1.0 respectively. These are highly comparable to the 11,874 genes in the *B. terrestris* gene set (v. 1.0), but lower than the 12,728 genes in the *B. impatiens* gene set (v. 2.1). Gene annotation of the *Bombus* assemblies presented in (Heraghty et al., 2020; Sun et al., 2020) also resulted in a greater number of genes annotated, ranging from 13,325 to 16,970 genes. BUSCO analyses of the annotated genes in our assemblies do suggest that we are missing some genes from our annotations (complete BUSCO genes in BBAL_01 and BSYL_01 respectively: 86.6% and 86.9%). Repeat content of our assemblies is comparable to that of other *Bombus* assemblies, with 17.11% of BBAL_1.0 and 17.85% of BSYL_1.0 identified as repetitive by RepeatMasker (full details of repeat classes in supplementary table S3). These proportions are comparable to the repeat content inferred in the *B. terrestris* and *B. impatiens* genomes.

### Population resequencing indicates three species clusters, with little population structure within species

The locations of sample collections, which comprise 580 samples from across seven localities in the Rocky Mountains, CO, USA, are presented in Fig. 1 (full details in table S1, S2). Illumina sequencing resulted in an average genome coverage of 13.1x per sample, with 4.5 and 3.5 million SNPs identified in *B. balteatus* and *B. sylvicola* samples respectively (supplementary table S4). Neighbor-joining trees produced from samples of both groups of samples indicate that the *B. balteatus* samples form a single cluster whereas the *B. sylvicola* samples form two clusters (Fig 1). We previously inferred the largest cluster to be comprised of *B. sylvicola* samples whereas the second cluster is comprised of the cryptic species “incognitus” (Christmas et al., 2021). Among samples originally designated as *B. sylvicola*, 24% (n=67) are identified as “incognitus” whereas the remaining 76% (n=214) are *B. sylvicola. Bombus balteatus* (n=299) was the most commonly observed species across all localities. The proportions vary between localities (table S1). “Incognitus” samples were collected on six out of the seven mountains, whereas *B. balteatus* and *B. sylvicola* were found on all seven.

**Figure 1.**
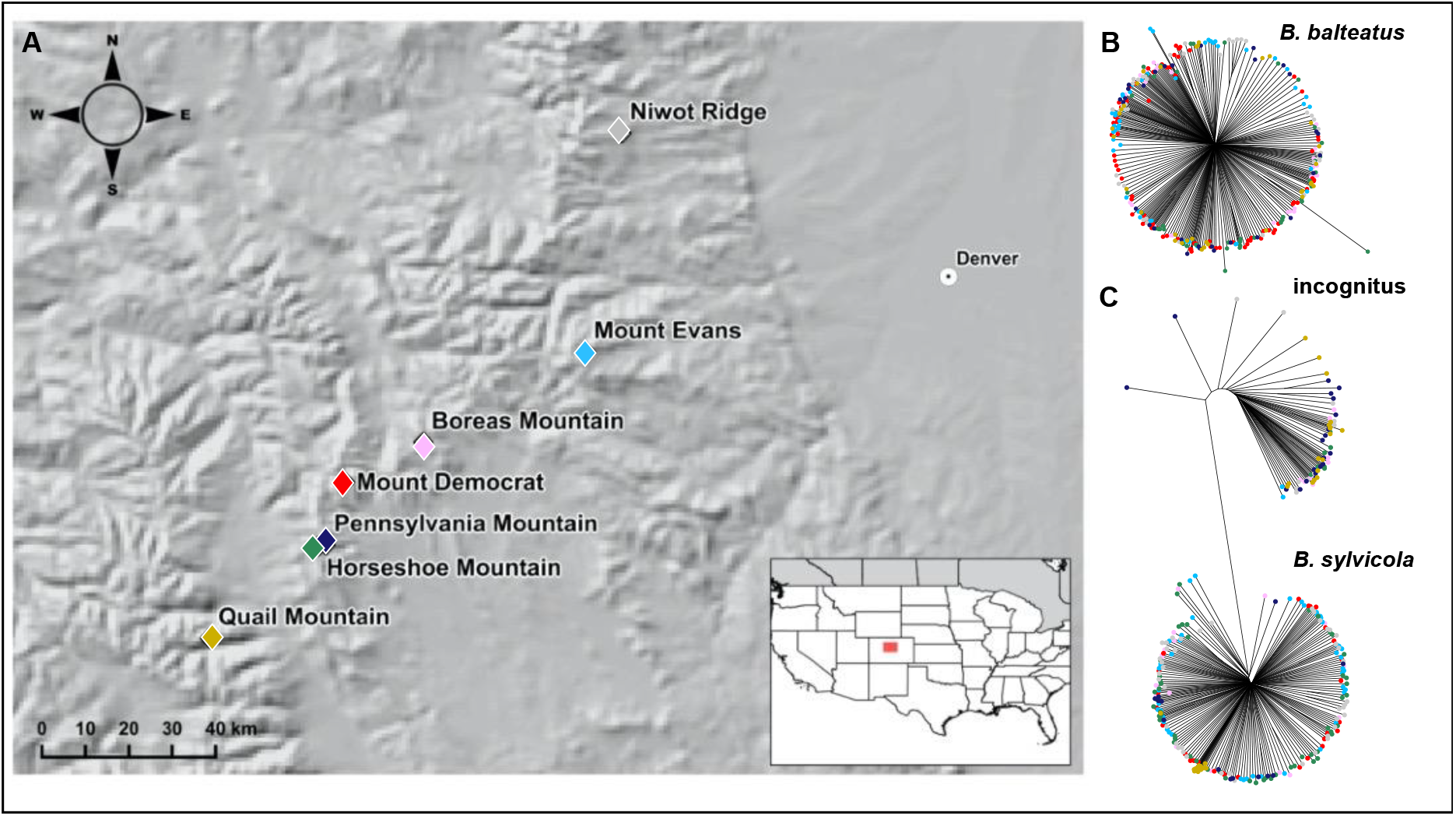
Population sampling and genetic variation in three *Bombus* species. (A) Map showing the sampling locations of *Bombus balteatus, B. sylvicola*, and “incognitus” on seven mountains in the Rocky Mountains, CO, USA. Insert indicates location in the USA. Neighbour-joining trees of (B) *B. balteatus* and (C) *B. sylvicola* and “incognitus” populations sampled across the seven mountains, based on genome-wide single nucleotide polymorphisms. Coloured tips indicate the mountain each sample was collected on, with colours corresponding to those in (A).

We calculated a pairwise kinship coefficient between samples to infer degrees of relatedness (Danecek et al., 2011; Manichaikul et al., 2010). We inferred a set of samples that were likely to be nestmates (sisters) based on them having the same maternal and paternal parents, under the realistic assumption that each colony is headed by a single monandrous female queen (Estoup et al., 1995). Those identified as sisters were always collected foraging on the same mountain and at the same or immediately neighboring site (table S1). This suggests that foraging bees were generally found in the immediate vicinity of their nests. Among the 299 *B. balteatus* samples, we identified 43 samples from 17 colonies that had another nestmate in the dataset, indicating that we sampled a total of 273 colonies. For the 214 *B. sylvicola*, we identified 14 samples from 7 colonies that met this criterion, indicating a total of 207 colonies. For 67 “incognitus”, we identified two samples from the same colony, indicating that we sampled 66 colonies.

We investigated the presence of population genetic structure within each species using several methods. Neighbor-joining trees (Fig. 1) do not reveal the presence of any substructure related to geography for *B. balteatus* or “incognitus”. For *B. sylvicola*, all of the samples from Quail Mountain (the furthermost southwestern site) cluster together, but do not separate from the rest on the tree. Principal component analysis further reflected the extremely low level of geographical structure across all species (Fig. 2). One slight exception to this trend is the Quail Mountain population of *B. sylvicola* which, shows a slight degree of separation from the other samples at the positive extreme of PC1 (Fig. 2B), although PC1 explains only 1.17% of the variance in the data. Outlier clusters seen in the PCA plots of *B. balteatus* and “incognitus” (Fig. 2A,C, circled) are composed of highly related individuals as shown in the kinship analysis, suggesting that the effect of kinship overrides any signal of genetic structure. After removing one of each pair of sisters, samples of the same species cluster tightly together and very little variance is explained by PC1 (Fig S1).

**Figure 2.**
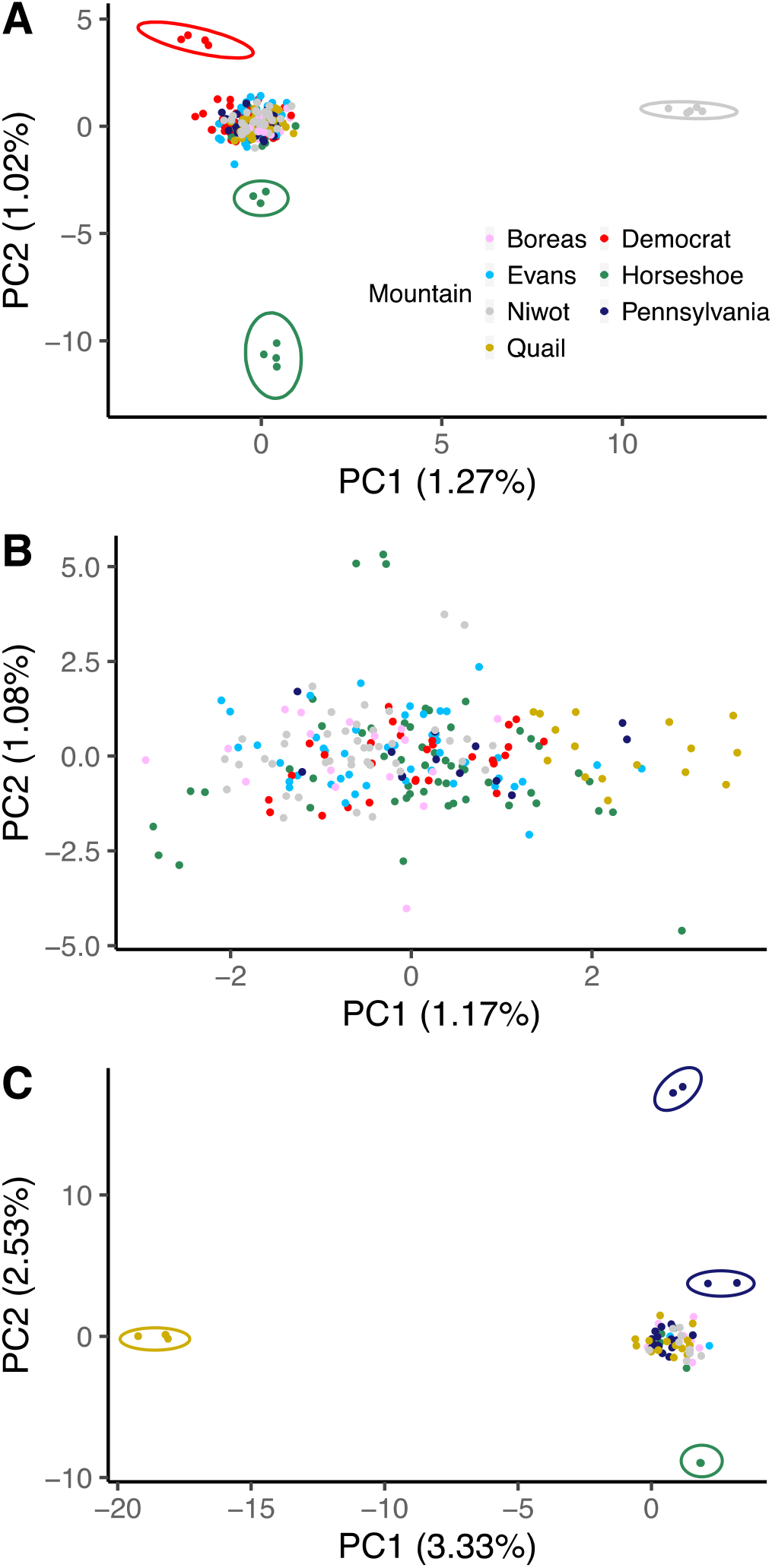
Principal component analysis of Rocky Mountain populations of three *Bombus* species. Plots show the first two principal components from principal component analyses of (A) *Bombus balteatus*, (B) *B. sylvicola* and (C) “incognitus” populations. Numbers in brackets on axes indicate the percentage variance explained by each principal component.

We assessed connectivity between colonies on different mountains by analyzing the association between kinship and geography using Mantel tests after removing sisters from the datasets (Fig. 3). We found no significant relationship between geographic distance and relatedness for *B. balteatus* (Mantel R = −0.03 (95% ci: −0.04 - −0.02), p value = 0.11; Fig. 3A,D). There were small but significant negative relationships for *B. sylvicola* (Mantel R = - 0.06 (95% c.i. −0.07 - −0.04), p value = 0.0001; Fig. 3B,E) and “incognitus” (Mantel R = −0.04 (95% ci: −0.06 - −0.03), p value = 0.029; Fig. 3C,F), demonstrating greater relatedness at closer locations. In these species, Mantel correlograms indicate significant associations between kinship and geographical distance only at shorter distances (from the same or neighboring mountains), which are no longer significant at greater distances (Fig. 3E-F). These results indicate that the limited population structure is mainly driven by the occurrence of a few highly-related samples from the same localities in the dataset.

**Figure 3.**
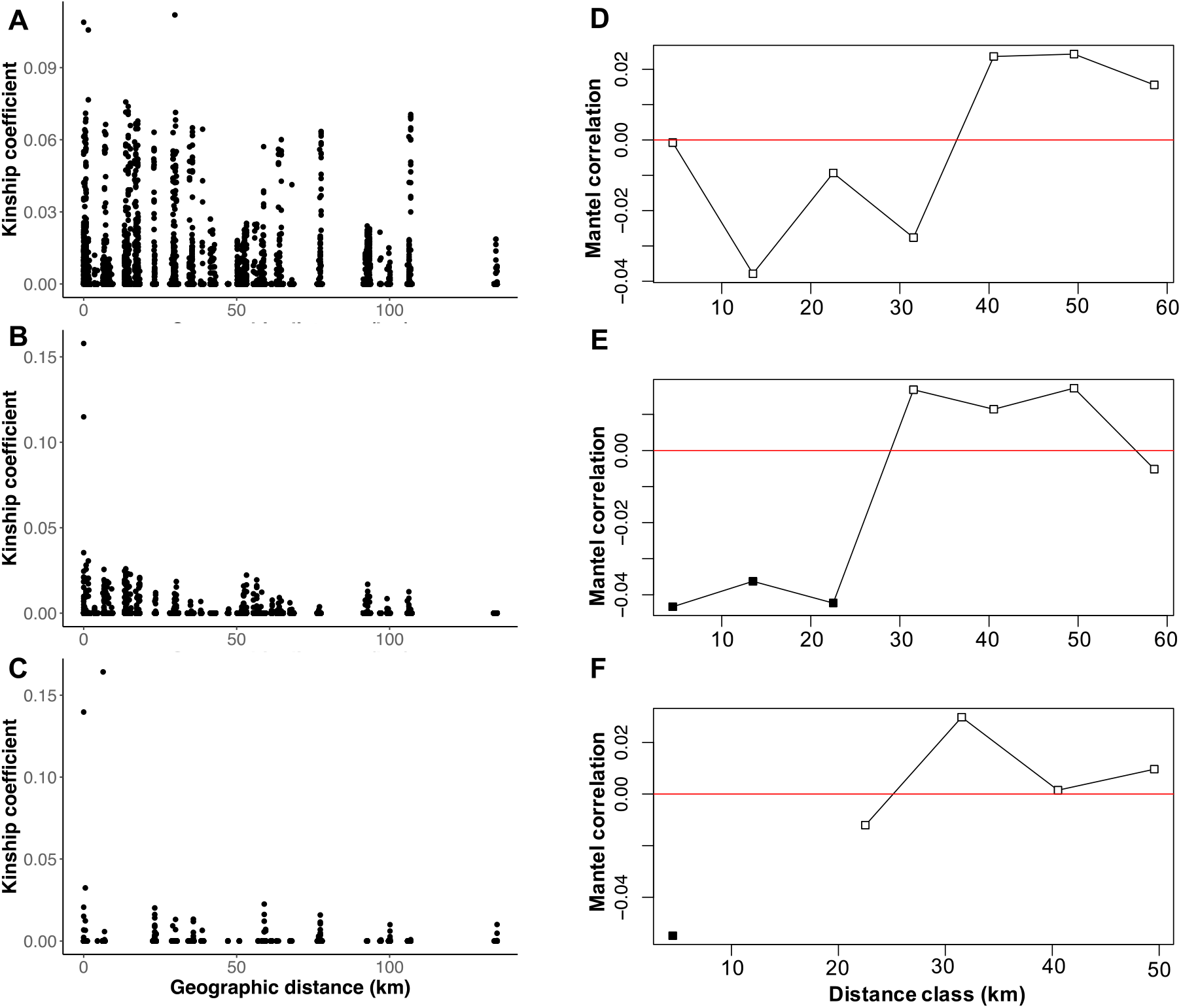
Correlations between kinship and geographic distance. Geographic distance between all pairs of samples plotted against kinship coefficient (a measure of relatedness) between pairs are shown for (A) *Bombus balteatus*, (B) *B. sylvicola*, and (C) “incognitus”, where each point represents a pair. (D-F) Mantel correlograms showing the Mantel correlation between kinship and geographic distance over 10 km distance classes for (D) *Bombus balteatus*, (E) *B. sylvicola*, and (F) “incognitus”. Filled in boxes indicate significant correlations (p < 0.01). There were insufficient data points for calculating Mantel correlations at distance classes > 60 km for *B. balteatus* and *B. sylvicola*. For “incognitus”, distance class 10-20 km as well as all distance classes > 50 km lacked sufficient data.

### Decrease in tongue length is ongoing in all species

We made morphological measurements for each sample (table S2). Intertegular distance is considered a proxy for body size (Cane, 1987). We also measured the length of two parts of the tongue: glossa and prementum. These two tongue measures are correlated in all three species (*B. balteatus*, Pearson’s r = 0.45, p < 0.001; *B. sylvicola*, Pearson’s r = 0.38, p < 0.001; incognitus, Pearson’s r = 0.53, p < 0.001, Figs. S2-4). Only glossa is considered further, as it was measured by Miller-Struttmann et al. (2015) and is the tongue part with most relevance to feeding (Cariveau et al., 2016). Glossa is also significantly correlated with intertegular distance (*B. balteatus*, Pearson’s r = 0.41, p < 0.001; *B. sylvicola*, Pearson’s r = 0.26, p < 0.001; incognitus, Pearson’s r = 0.56, p < 0.001, Figs. S2-4).

A comparison of phenotypes between species shows that glossa length is substantially longer in *B. balteatus*, consistent with previous knowledge (ls-means = 4.72 mm, 3.34 mm, and 3.40 mm for *B. balteatus, B. sylvicola* and “incognitus” respectively; Tukey test, t ratio = 12.25 and 21.17, p < 0.001 for *B. balteatus* compared to *B. sylvicola* and “incognitus”; Fig. 4A). Average glossa length is extremely similar in *B. sylvicola* and “incognitus” (Tukey test, t ratio =0.50, p = 0.87), suggesting that this character does not distinguish this newly-discovered species from *B. sylvicola. Bombus balteatus* bees are significantly larger bees as indicated by a greater intertegular distance, and *B. sylvicola* is significantly larger than “incognitus” (mean intertegular distance = 4.00 mm, 3.83 mm, 3.58 mm for *B. balteatus, B. sylvicola* and “incognitus” respectively; ANOVA, F=28.75, p < 0.001, Tukey post-hoc test revealed differences are significant among all species at p < 0.001; Fig. S5A).

**Figure 4.**
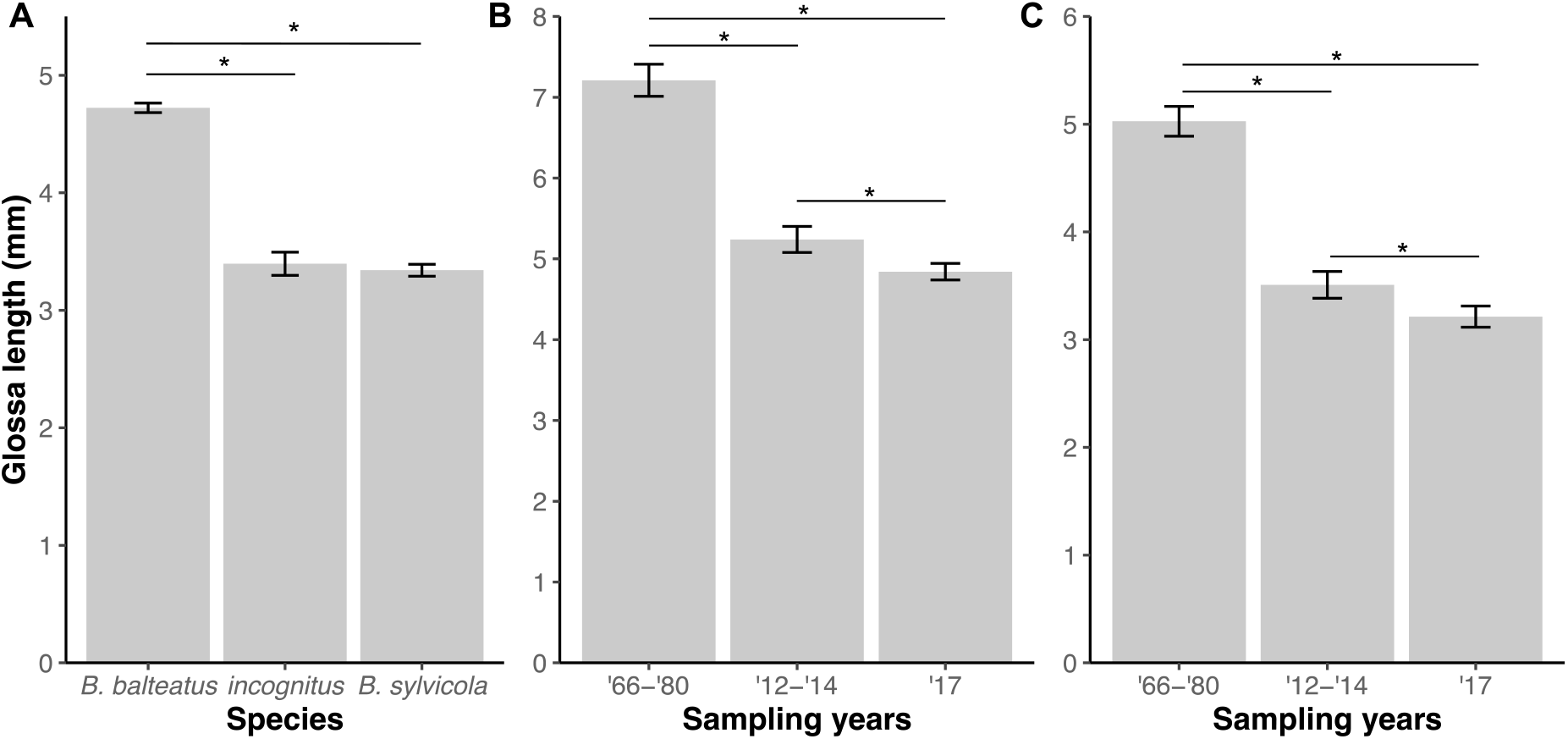
Comparisons of glossa lengths between species, years, and locations. (A) Least-squares mean (ls-mean) glossa lengths correcting for body size for the three *Bombus* species, based on measurements of the 2017 collections. Least-squares mean glossa lengths, correcting for body size and sample location, per sampling period for (B) *B. balteatus* and (C) *B. sylvicola* and “incognitus” combined. In all plots, error bars represent ls-means ± standard error and significant differences between classes are indicated by asterisks (p < 0.05).

We next compared the distribution of glossa lengths in our samples with historical data. We combined *B. sylvicola* and “incognitus” for this analysis because they do not differ in glossa length on average and they were not distinguished in previous studies. We find an overall trend of decreasing glossa length in both *B. sylvicola*-incognitus and *B. balteatus* over time (Fig. 4B,C). This was reported previously between the collections from 1966-1980 and from 2012-2014 for both species (Miller-Struttmann et al., 2015). Here we show that this trend is continuing, with further reductions in glossa length between the 2012-2014 collections and our collections from 2017. These reductions are significant for both *B. sylvicola*-incognitus (ls-means: 2012-2014 = 3.51 mm, 2017 = 3.21 mm; Tukey test, t-ratio = 3.198, p = 0.004) and *B. balteatus* (ls-means: 2012-2014 = 5.24 mm, 2017 = 4.84 mm; Tukey test, t-ratio = 2.79, p = 0.015). In contrast, there is no significant trend shown by intertegular distance (Fig. S5B,C), although for *B. sylvicola*-incognitus, intertegular distance is significantly larger in both the 1966-80 and 2017 samples compared to the 2012-2014 samples (ANOVA, F = 15.28, p < 0.001. Tukey post-hoc test, p < 0.001).

For samples collected in 2017, we compared the distribution of glossa lengths for each species among the seven mountains where they were collected (Fig. S6). There is an overall trend for glossa lengths to be longer in bees from Mount Evans than on other mountains in all species. The distributions for *B. balteatus* and *B. sylvicola* show significant differences, with samples from Mount Evans having significantly longer glossae than on Mount Democrat, Horseshoe Mountain, and Pennsylvania Mountain in *B. balteatus* (ls-mean glossa length on Mt. Evans = 5.34 mm, other mountains 4.51 – 4.73 mm; Tukey p < 0.01 in each case; Fig. S6A) and significantly longer glossae than on all other mountains except Niwot Ridge in *B. sylvicola* (ls-mean glossa length on Mt. Evans = 3.77 mm, other mountains 3.02 – 3.24 mm; Tukey p < 0.01 in each case; Fig. S6B). A similar but non-significant trend could also be seen for “incognitus” (Fig. S6C). Comparisons of intertegular distance between mountains revealed that *B. balteatus* bees sampled on Niwot Ridge are significantly larger than those on Boreas Mountain, Mount Democrat, Horseshoe Mountain, and Quail Mountain (Niwot Ridge mean intertegular distance = 4.25 mm, other mountains 3.84 – 4.00 mm, ANOVA, F = 4.09, p < 0.001, Tukey post-hoc test p < 0.05 in four comparisons; Fig. S7A). We did not identify any significant differences in intertegular distance of bees across mountains for *B. sylvicola* (ANOVA, F = 1.83, p = 0.10) or “incognitus” (ANOVA, F = 1.62, p = 0.17; Fig. S7B,C).

### Genome-wide association study identifies a SNP associated with a proxy for body size

We used a linear mixed-model analysis to search for correlations between allele frequencies and quantitative trait variation in *B. sylvicola* and *B. balteatus*. We used the traits glossa length, intertegular distance and glossa length corrected for intertegular distance (see Methods). One from each pair of sisters identified in the kinship analysis was removed from all analyses to reduce the level of relatedness in the sample set. We detected a genome-wide significant association between intertegular distance and a SNP in intron 9 of the *CTRB1* (Chymotrypsinogen B1) gene on chromosome 10 in *B. sylvicola* (p < 0.001 after Bonferroni correction; Fig. 5). The SNP is found at an allele frequency of 0.14 across all populations, with 41 individuals heterozygous and nine homozygous for the allele. The allele is present on all mountains, ranging in allele frequency from 0.06 (Quail Mountain) to 0.24 (Niwot Ridge). The *CTRB1* gene is involved in production of chymotrypsin, a proteolytic digestive enzyme (Burgess, Malone, & Christeller, 1996; Giebel, Zwilling, & Pfleiderer, 1971; Grogan & Hunt, 1979; Lazarević & Janković-Tomanić, 2015; Matsuoka et al., 2015; Rawlings & Barrett, 1994). Intron 9 is a large 18 Kbp intron and the significant SNP is located 2 Kbp upstream of exon 10. The GWAS significance suggests it is of likely functional importance and its location is suggestive of a regulatory element, such as an intron splice enhancer or silencer.

**Figure 5.**
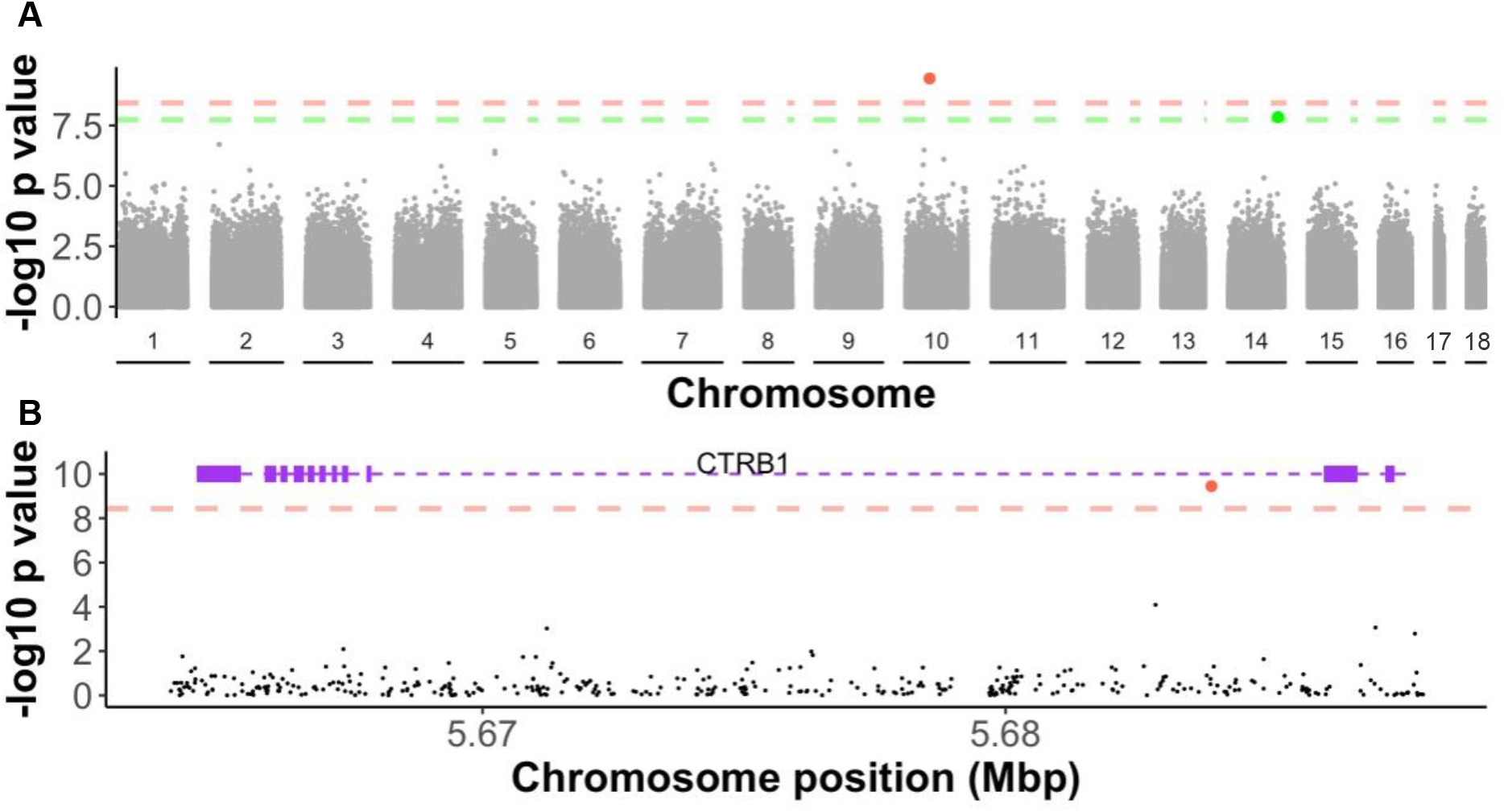
Genome-wide association Manhattan plot. (A) The -log_10_ p-values from an association test using a mixed linear model between allele frequencies and intertegular distance (ID) are plotted for 2,705,361 genome-wide SNPs against their genome positions over the 18 *Bombus balteatus* pseudochromosomes. Red dashed line indicates genome-wide significance after Bonferroni correction for p = 0.01. A single SNP on chromosome 10, coloured and circled in red, had a -log_10_ p-value above this threshold. Green dashed line indicates genome-wide significance after Bonferroni correction for p = 0.05. Two neighbouring SNPs located on chromosome 14, coloured green, had -log_10_ p-values above this threshold. (B) Zoomed-in plot of the significant SNP in chromosome 10 showing its location within an intron of the *CTRB1* gene. Exons of this gene are represented by purple boxes, introns are represented by dashed purple lines.

Two further SNPs, located 4 bp apart on chromosome 11, also significantly associated with intertegular distance (p = 0.039 after Bonferroni correction). However, they are not located in or close to any genes according to our annotation, so any potential functional significance of these SNPs is unknown. We did not find any significant associations between allele frequencies and intertegular distance in *B. balteatus* (Fig. S8). Furthermore, no significant associations were found between allele frequencies and glossa length, or glossa length corrected for intertegular distance in either species (Fig. S8). This indicates that the trait has a polygenic or mainly environmental component. A power analysis (Visscher et al., 2014) revealed that our datasets have sufficient power (>0.8) to detect a QTL with a h^2^ of ~0.1 but that loci with lower effect sizes would likely be undetectable.

### Limited evidence that morphological variation has a genetic component

We performed GCTA-GREML analyses (Yang et al., 2011) to determine the genetic component of trait variation for *B. sylvicola* and *B. balteatus* (Fig. 6, supplementary table S5). The “incognitus” samples were not included here due to smaller sample size. When each species’ entire dataset is considered, we detect a heritable component for both glossa length and intertegular distance in both species, where the 95% confidence interval does not overlap zero. For *B. sylvicola*, but not *B. balteatus*, this is also the case when intertegular distance is introduced as a covariate to account for body size when considering glossa length. However, when we remove samples that we infer to be workers from the same colony from the kinship analysis, we only detect a heritable component with 95% confidence for intertegular distance in *B. balteatus* (all others then overlap with zero). This indicates that a large proportion of the signal of covariance between genetic and trait variance is due to the presence of sisters from the same colony. This could be a confounding factor because environmental effects such as level of nutrition and colony health are likely to covary with colony. However, such covariation could also reflect genetics. The level of dependency of phenotype on genotype is therefore unclear from this analysis.

**Figure 6.**
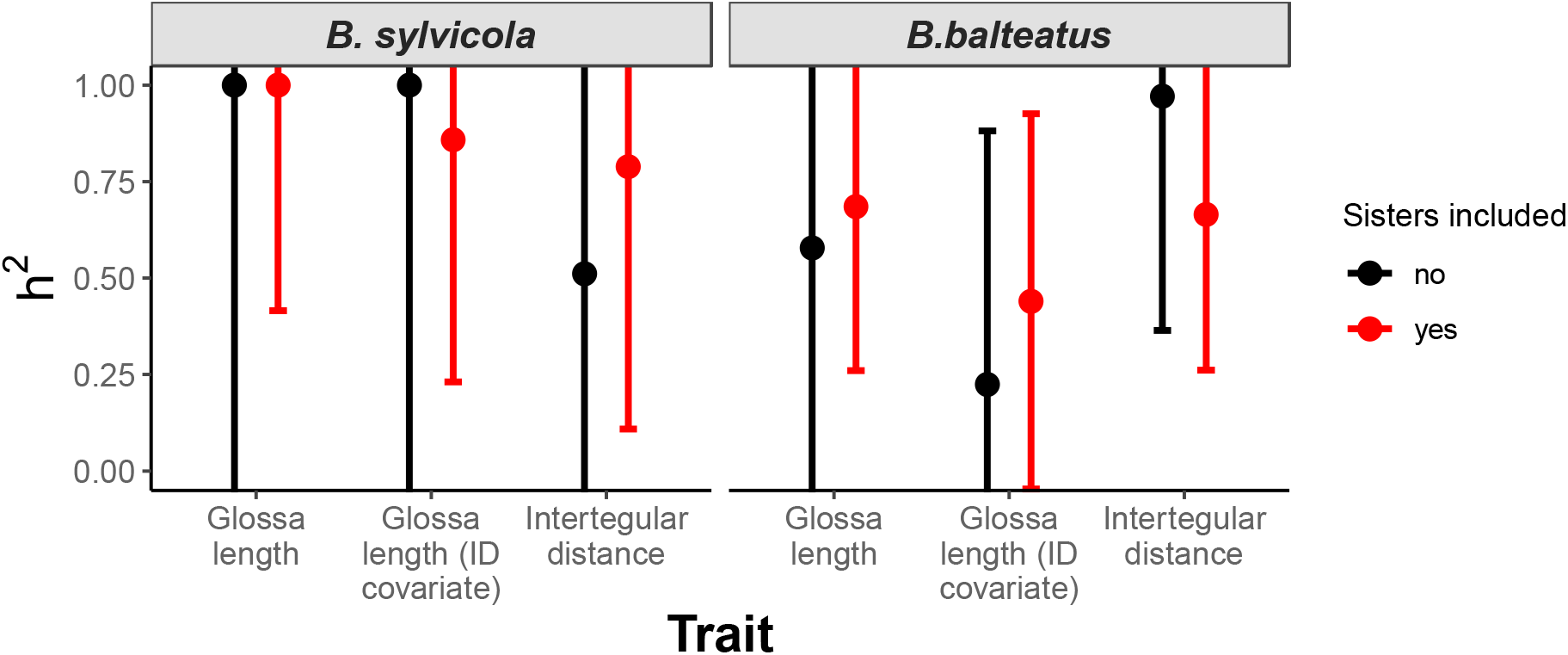
Estimates of SNP-heritability for two traits in *Bombus* species. The proportion of variance in glossa length and intertegular distance (ID) explained by all genome-wide SNPs (the SNP-based heritability) was calculated using the genome-base restricted maximum likelihood (GREML) method in GCTA. These were calculated separately for *B. sylvicola* (left panel) and *B. balteatus* (right panel). We included intertegular distance as a covariate with glossa length to control for body size. Error bars indicate 95% confidence intervals around estimates. Each analysis was carried out twice, once with all samples (red), and a second time with one from each pair of sisters removed (black), to assess the effect of including highly related samples in the analysis.

The 95% confidence intervals for the estimate of heritability have a wide span and overlap zero in many cases. The power to determine heritability accurately depends on several factors, including sample size, the structure of relatedness in the samples, and the true value of heritability. When samples are all completely unrelated, it has been estimated that > 3,000 samples are required to achieve a SE below 0.1 (Visscher et al., 2014). For datasets of unrelated samples of the size we have analyzed here, we estimate a power of ~0.06, or ~6% probability, to detect h^2^ > 0 for a trait with a SNP-heritability of 0.5 and a type 1 error rate (α) of 0.05. Greater sample size or higher levels of relatedness within the dataset would therefore increase accuracy of our heritability estimates.

## DISCUSSION

We generated a high-quality reference genome and performed genome sequencing of a population sample of the bumblebee species *B. balteatus*. Combining these data with a previous dataset from *B. sylvicola* produced the most extensive population genomics dataset for bumblebees to date, with 580 samples from these two species and the newly-discovered “incognitus” collected from high-altitude locations in Colorado, USA. These data were combined with morphometric measurements from each sample and used to investigate genetic structure, morphological evolution and the genetic basis of phenotypic traits in these populations. Our main findings are 1) no evidence for gene flow between species, including the recently-inferred “incognitus”, but lack of population structure within species, indicating high levels of connectivity among subpopulations. 2) An ongoing decrease in tongue (glossa) length compared to historical measurements, previously inferred to reflect the effects of decreasing floral abundance on bumble bee foraging (Miller-Struttmann et al., 2015). 3) An association between body size and variation at a single SNP located in a gene related to production of a digestive enzyme in *B. sylvicola*. 4) A lack of conclusive evidence for a genetic component to tongue length in both species. Our analysis is unable to determine the degree of heritability underlying tongue length or body size but is consistent with these traits having a polygenic genetic component.

This investigation adds to the number of bumblebee species with reference genomes available. The genomes of the common species *B. impatiens* and *B. terrestris* were sequenced using short-read technologies (Sadd et al., 2015). Genome assemblies are also available for representatives of all 15 *Bombus* subgenera (Sun et al., 2020) based on short-read sequencing of jumping libraries and scaffolding using Hi-C, resulting in contig N50 of 325 kb. In addition to the two genomes produced here, the genomes of the North American species *B. terricola, B. bifarius, B. vancouverensis*, and *B. vosnesenskii* have also been produced using long-read sequencing (PacBio or ONT) and have megabase-scale contig N50s (Heraghty et al., 2020; Kent et al., 2018).

Identification of the previously-unknown species “incognitus” demonstrates the power of population-scale genome sequencing to discover new cryptic variation (Christmas et al., 2021). None of the morphological features that we studied could distinguish between *B. sylvicola* and “incognitus”. *Bombus sylvicola* did however have a significant tendency to be collected at higher altitudes and have larger body size. The wider distribution of “incognitus” is unknown, as is the true composition of populations described as *B. sylvicola* in other geographical locations. Existing collections of *B. sylvicola* from Colorado are likely to be a mixture of both species. However, importantly for this study, there were no significant differences in glossa or prementum length between the two species.

We find no evidence for population structure among samples of any of the three species studied here. This indicates an absence of long-term geographical barriers to gene flow among the high-elevation localities included here. Furthermore, although we observe elevated degrees of relatedness among foragers collected on the same mountain, there is no significant tendency for elevated relatedness of foragers caught on nearby mountains, indicating a lack of population structure at this scale. Foragers inferred to belong to the same nest (i.e. with identical parents) were always found in the same or adjacent sites, consistent with the limited foraging distance (on average < 110m) inferred previously by Geib et al. (2015) for these species.

Studies of several bumblebee species across continental USA, including *B. bifarius* and *B. vosnesenkii*, also found in Colorado and across western North America, indicate weak geographical differentiation even at distances over 1000 km (Ghisbain et al., 2020; Lozier et al., 2011). However, the complex topography of mountain ranges likely modulates gene flow among populations (Lozier et al., 2013, 2011). Populations of *B. bifarius* are inferred to be more fragmented in the southern portions of their ranges, where they occur at higher elevations (Jackson et al., 2018). The species studied here, *B. sylvicola* and *B. balteatus*, are generally restricted to high-elevation localities above the tree line separated by forest (Williams, Thorp, Richardson, & Colla, 2014). However, our analysis indicates that this complex landscape has not historically limited mating and dispersal between localities in the same mountain range. This is consistent with previous studies of these species that found substantial genetic differentiation between mountain ranges in the Pacific Northwest, but not within the same mountain range (Koch et al., 2017; Whitley, 2018).

Morphological change over time has already been documented in the species studied here, with a significant reduction in tongue length observed since 1966 (Miller-Struttmann et al., 2015). During this time period, there has also been a shift in species composition, with shorter-tongued species becoming more predominant. Previous collections of *B. sylvicola* in this region are likely a mixture of *B. sylvicola* and “incognitus”, as the existence of “incognitus” was unknown. However, as *B. sylvicola* and “incognitus” do not differ in glossa length, undetected changes in the relative abundance of these species over time are unlikely to be responsible for the observed shifts in glossa length observed in samples that are defined as *B. sylvicola*. This effect could, however, potentially contribute to the observed variation in body size over time as *B. sylvicola* have a significantly greater intertegular distance on average than “incognitus” (Fig. S5).

A major finding that we present here is that the temporal shift toward shorter tongue length is ongoing. In addition to the significant reduction previously observed over a period of six decades up until 2014, we also observe a significant reduction over a much shorter time period between 2012-14 and 2017 in both *B. balteatus* and *B. sylvicola*-incognitus. Mean tongue length reportedly decreased on average by 0.61% annually in a period between 1966 and 2014 for *B. sylvicola* (Miller-Struttmann et al., 2015). According to our recent measurements, the corresponding decrease in 3-5 subsequent years has been 12.9 %, or 3.2% per year (compared to the 2012-14 average).

Decreases in tongue-length have been previously attributed to an advantage of shorter tongues for foraging on a wide range of flowers. Such generalist foraging could be favored due to a general decline of floral resources observed in connection with warmer summers (Miller-Struttmann et al., 2015). While both long- and short-tongued bees are capable of accessing nectar in plants with short flower tubes, short-tongued bees are more efficient at foraging from them (Inouye, 1980) and are also more efficient at foraging across a diversity of flower tube depths (Arbulo, Santos, Salvarrey, & Invernizzi, 2011; Geib & Galen, 2012; Plowright & Plowright, 1997). Short-tongued bumble bees often forage from a wider suite of plants both in these alpine bumble bee species (Miller-Struttmann et al., 2015) and others (Goulson & Darvill, 2004; Goulson et al., 2008b; Heinrich, 2004; Huang et al., 2015). These considerations indicate that shorter tongues are likely advantageous for generalist foraging. Although short-tongued bees are unable to pollinate plants with long flower tubes, they are likely important for supporting a diverse array of flowering plants.

It is unclear whether the tongue-length changes we observe are the result of natural selection or phenotypic plasticity. There are now many examples of rapid evolution of traits driven by natural selection due to climatic events in the wild, such as a shift in beak shape over a period of a few years in Darwin’s finches related to competition for food during drought (Lamichhaney et al., 2016) and genomic shifts related to cold-tolerance in the green anole lizard related to severe winter storms (Campbell-Staton et al., 2017). Human activity may also promote rapid adaptation by natural selection, as demonstrated by the evolution of longer beaks in great tits promoted by the use of bird feeders in the UK (Bosse et al., 2017). Global climate change is also expected to be a driver of rapid evolution, but it is often difficult to establish whether trends of phenotypic change result from evolution or phenotypic plasticity (Merilä & Hendry, 2014). Long-term changes in breeding timing in great tits (Charmantier et al., 2008) and body mass in red-billed gulls (Teplitsky, Mills, Alho, Yarrall, & Merilä, 2008) that track temperature increases have been shown to result solely from phenotypic plasticity.

To address the causes of the changes in bumblebee tongue length observed here, we attempted to uncover the genetic basis of morphological variation using genome-wide association studies (GWAS) and genome-wide complex trait analysis (GCTA). No significant associations were observed for tongue length in the genome of any of the species using GWAS. However, a significant association with body size was identified in *B. sylvicola* at a single SNP in the *CTRB1* (Chymotrypsinogen B1) gene. The lack of association at neighboring SNPs could be attributable to high rates of meiotic recombination and low levels of linkage disequilibrium in bumblebees (Kawakami et al., 2019). The *CTRB1* gene produces a precursor of chymotrypsin, a key digestive enzyme in animals, which is likely important for pollen digestion in bees (Burgess et al., 1996; Giebel et al., 1971; Grogan & Hunt, 1979; Lazarević & Janković-Tomanić, 2015; Matsuoka et al., 2015; Rawlings & Barrett, 1994). The associated SNP is found in a large intron (18 Kbp) of this gene, 2 Kbp upstream of the nearest exon. We lack any functional annotation for this SNP but its location and GWAS significance suggests it may play a role in gene regulation. For example, if located within an intronic splicing enhancer or silencer, it may result in variation in expression levels of different isoforms of the CTRB1 enzyme being produced. Such changes could influence body size through an effect on the efficiency of digestion.

We detected a significant signal of covariance of trait measurements with relatedness for both tongue length and intertegular distance using GCTA when closely-related individuals were included in the dataset. However, no significant signal of heritability in morphological traits was detected when the dataset was purged of close relatives. The first result could potentially reflect a genetic component, but it is not possible to disentangle this component from the shared environmental effects due to close relatives sharing a nest. This analysis is therefore underpowered to detect genetic effects due to low sample size and a general lack of related individuals in the dataset. A possible explanation for our results is that the morphological traits tongue length and intertegular distance have a highly polygenic genetic component, with multiple QTL of small effect (h^2^ < 0.1), such that GWAS and GCTA were underpowered to detect QTL and estimate heritability. The size of the genetic component of variation in both traits is therefore still unknown in each species. These results contrast with findings regarding the genetic control of bumblebee coloration, in which variation at single loci control intraspecific variation in color of abdominal segments (Rahman, Cnaani, Kinch, Grishin, & Hines, 2021; Tian et al., 2019).

It therefore remains unclear whether the observed changes in tongue length are caused by natural selection acting on functional genetic variation, or whether they are the result of phenotypic plasticity. There is large variation in body size between individuals in bumblebee nests, indicating substantial phenotypic plasticity in this trait (Peat, Tucker, & Goulson, 2005). This variability could be explained by a correlation between larval position in the nest and size, which is related to feeding intensity (Couvillon & Dornhaus, 2009). There is evidence from solitary bees of an effect of temperature on body size (Scaven & Rafferty, 2013), although such an effect is likely to be less pronounced in social insects such as bumblebees that regulate the temperature of their nests. However, it is unclear whether tongue length also exhibits plasticity, as this trait correlates with body size among nestmates (Peat et al., 2005), indicating that plasticity in tongue length independent of body size is less pronounced. However, our observation of significant differences in mean tongue length between mountains could potentially reflect phenotypic plasticity.

In summary, we have shown that decreasing tongue length observed over the last six decades is still ongoing in populations of the bumblebee species *B. sylvicola* and *B. balteatus* (and the recently-reported “incognitus”) at high altitudes in Colorado. We have generated the genomic resources to study these species on the genetic level in the form of highly-contiguous genome assemblies and comprehensively characterized genome variation. Our results indicate that the complex topology of the mountain landscape in Colorado does not contain significant barriers to mating and dispersal of bumblebees in the study area. We are unable to conclude whether the observed morphological shifts over time are the result of genetic adaptation or phenotypic plasticity. Alpine ecosystems are particularly sensitive to climate change and may suffer its effects before other geographical regions (Elsen & Tingley, 2015). Continued monitoring of changes in morphology in these and other populations in these regions could be highly valuable for understanding how pollinators respond to the effects of a warming climate, the degree to which rapid evolution by natural selection or phenotypic plasticity are possible, and the effects of the changes on these ecosystems.

## DATA AND CODE AVAILABILITY

All sequence data presented here are available at NCBI under BioProject IDs PRJNA646847 and PRJNA704506. Scripts and annotation parameters used in the study are available at GitHub (https://github.com/matt-webster-lab/alpine_bumblebees). VCF files containing genotypic variation are available at the Dryad Digital Repository (https://doi.org/10.5061/dryad.mgqnk990p).

## ACKNOWLEDGEMENTS

This research was funded by Swedish Research Council Formas grant 2016-00535 and a SciLifeLab National Biodiversity Project NP00046 to MTW. MK is financially supported by the Knut and Alice Wallenberg Foundation as part of the National Bioinformatics Infrastructure Sweden at SciLifeLab. The authors acknowledge support from the National Genomics Infrastructure in Stockholm and Uppsala funded by Science for Life Laboratory, the Knut and Alice Wallenberg Foundation and the Swedish Research Council. The computations and data handling were enabled by resources provided by the Swedish National Infrastructure for Computing (SNIC) at UPPMAX partially funded by the Swedish Research Council through grant agreement no. 2018-05973. We thank the Mountain Research Station (U.C. Boulder), Mt. Evans Research Station (University of Denver), the U.S. Forest Service, Dr. Rosemond Greinetz and the Mountain Area Land Trust for providing assistance and access to field sites. We thank Jacqueline Staab and Michael Osbourn who aided in specimen collection, identification and preservation and Lisa Danback, Kate Schleicher, Ying Thao, Anna Grobelny and Breana Cook who carried out phenotypic measurements.

## AUTHOR CONTRIBUTIONS

Designed research: MJC, NEM-S, JCG, MTW; Performed research: all authors; Analyzed data: MJC, JCJ, OW, IB, MK, MTW; Wrote the paper: MJC, MTW.

