## supplementary figures for "A genomic and morphometric analysis of alpine bumblebees: ongoing reductions in tongue length but no clear genetic component"

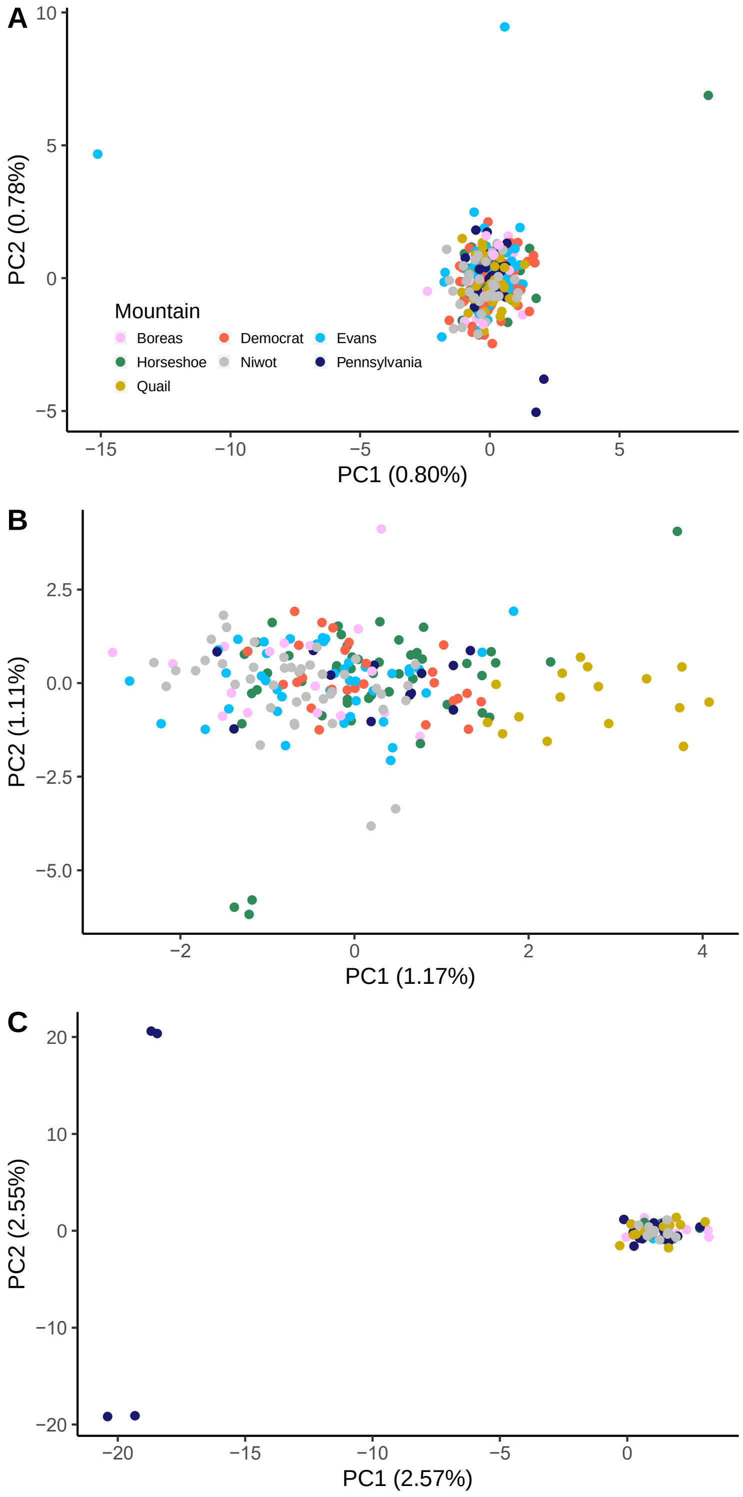


**Figure S1. Principal component analysis of Rocky Mountain populations of three *Bombus* species after removing one from each pair of sisters.** Plots show the first two principal components from principal component analyses of (A) *Bombus balteatus*, (B) *B. sylvicola* and (C) "incognitus" populations. Numbers in brackets on axes indicate the percentage variance explained by each principal component. The tight clustering and extremely low levels of variation explained by the PCs demonstrates that population genetic structure is minimal among the seven mountain sites. Outliers in (C) are two pairs of highly-related samples.

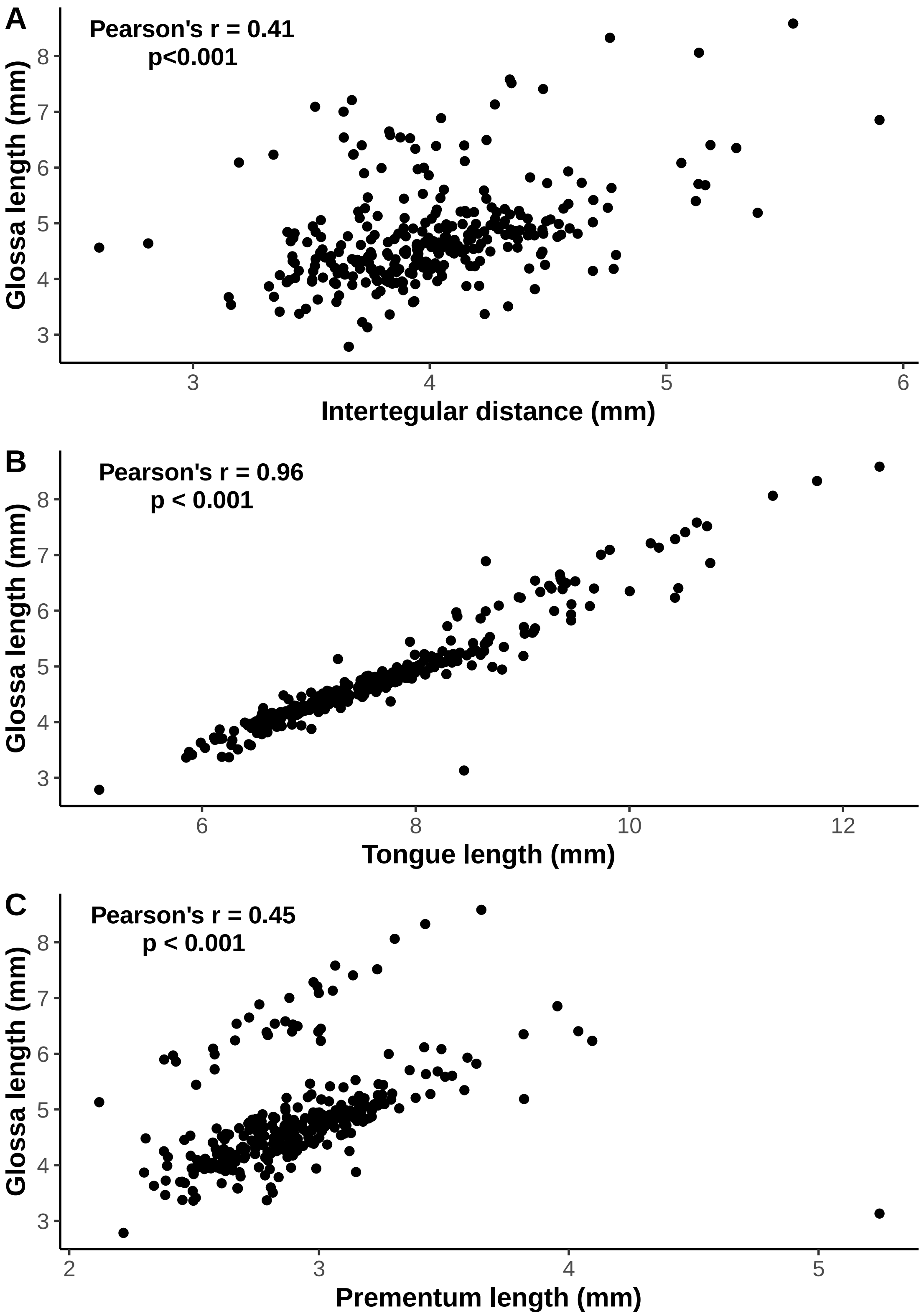


**Figure S2 Scatter plots of traits measured in *B. balteatus* samples collected in 2017.** (A) glossa length (mm) against intertegular distance (mm; n = 303). (B) glossa length (mm) against total tongue length (mm; glossa length + prementum length; n = 210). (C) glossa length (mm) against prementum length (mm; n = 210). Pearson's correlation coefficient (r) and p-values are indicated on each plot.

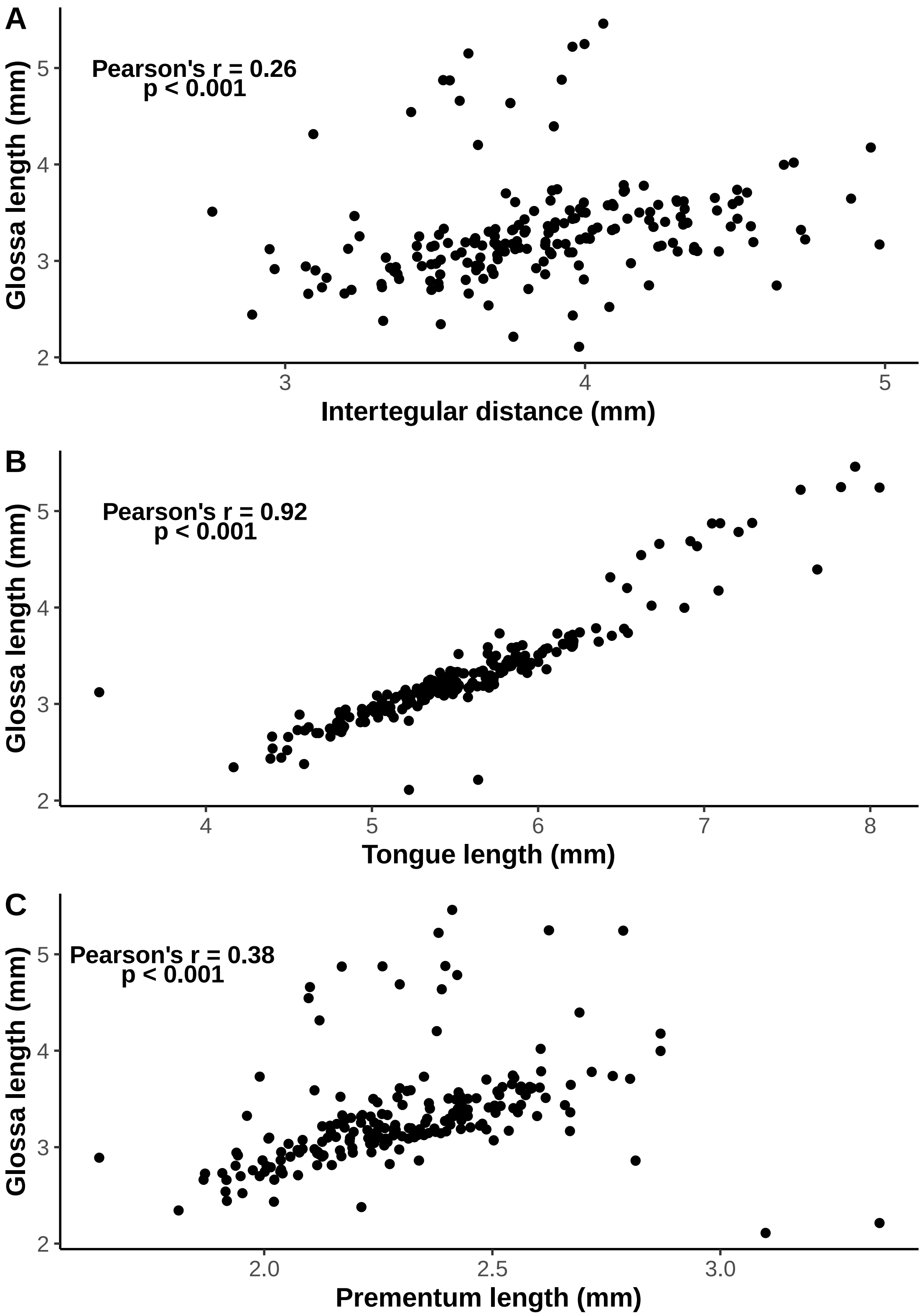


**Figure S3 Scatter plots of traits measured in *B. sylvicola* samples collected in 2017.** (A) glossa length (mm) against intertegular distance (mm; n = 189). (B) glossa length (mm) against total tongue length (mm; glossa length + prementum length; n = 205). (C) glossa length (mm) against prementum length (mm; n = 206). Pearson’s correlation coefficient (r) and p-values are indicated on each plot.

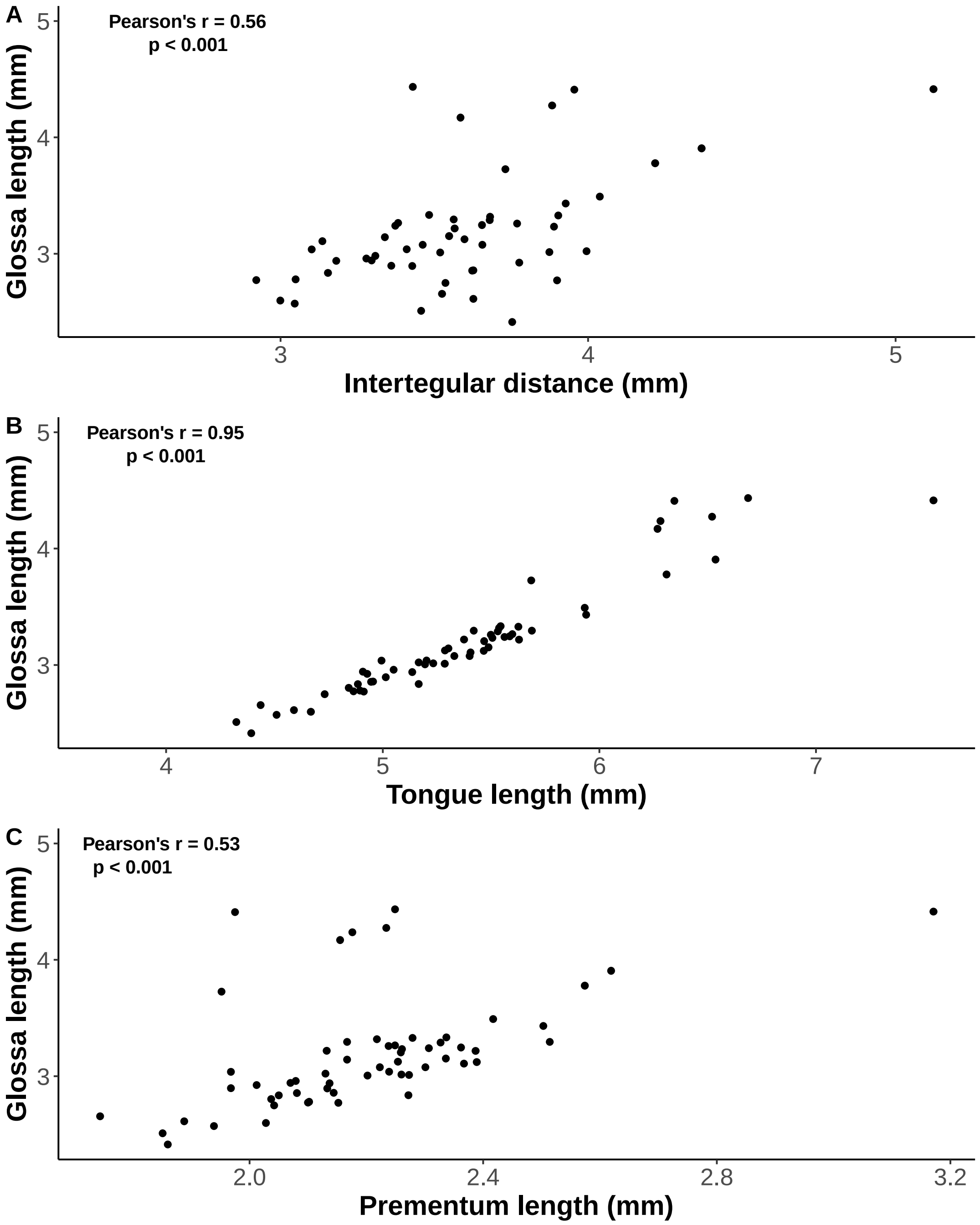


**Figure S4 Scatter plots of traits measured in incognitus samples collected in 2017.** (A) glossa length (mm) against intertegular distance (mm; n = 52). (B) glossa length (mm) against total tongue length (mm; glossa length + prementum length; n = 58). (C) glossa length (mm) against prementum length (mm; n = 59). Pearson's correlation coefficient (r) and p-values are indicated on each plot.

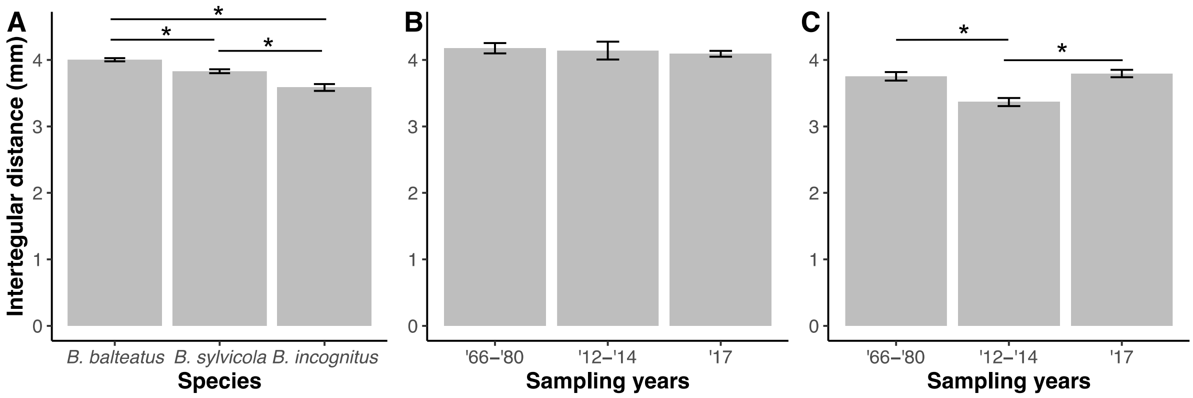


**Figure S5. Comparisons of intertegular distance between species and years.** (A) Intertegular distance for the three *Bombus* species, based on measurements of the 2017 collections. Intertegular distance per sampling period for (B) *B. balteatus* and (C) *B. sylvicola* and incognitus combined. In all plots, error bars represent ls-means ± standard error and significant differences between classes are indicated by asterisks ($p < 0.05$).

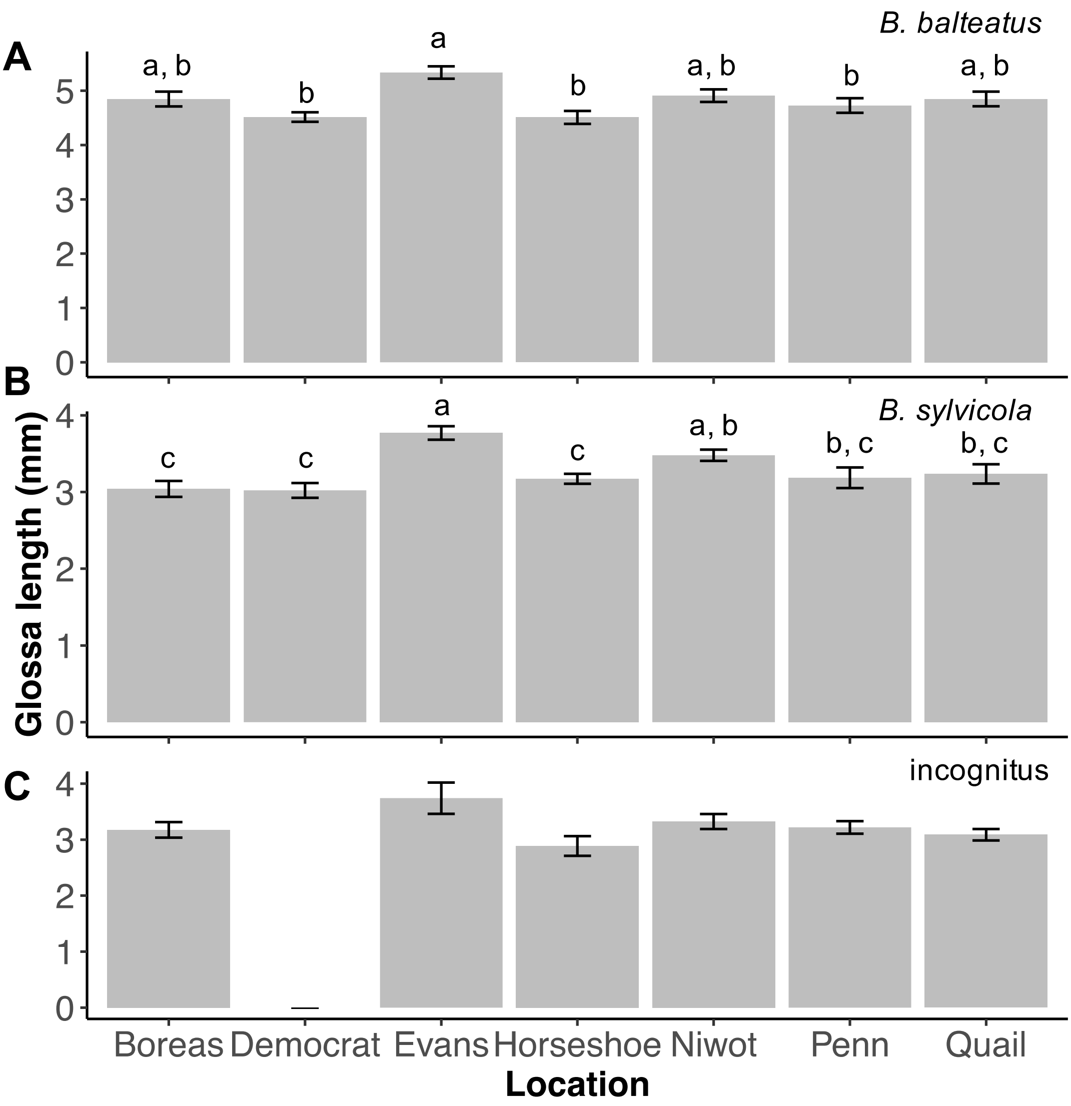


**Figure S6. Variation in glossa length across seven mountain locations in the Rocky Mountains for three *Bombus* species.** The least-square means (ls-means) of glossa length accounting for body size for (A) *Bombus balteatus*, (B) *B. sylvicola*, (C) incognitus. In each case, letters indicate significant differences between mountains based on Tukey's post hoc test, where locations not sharing letters are significantly different at $p < 0.05$. Error bars show ls-means $\pm$ standard error. Incognitus samples were not collected on Mount Democrat, and there were no significant differences in glossa lengths among mountains.

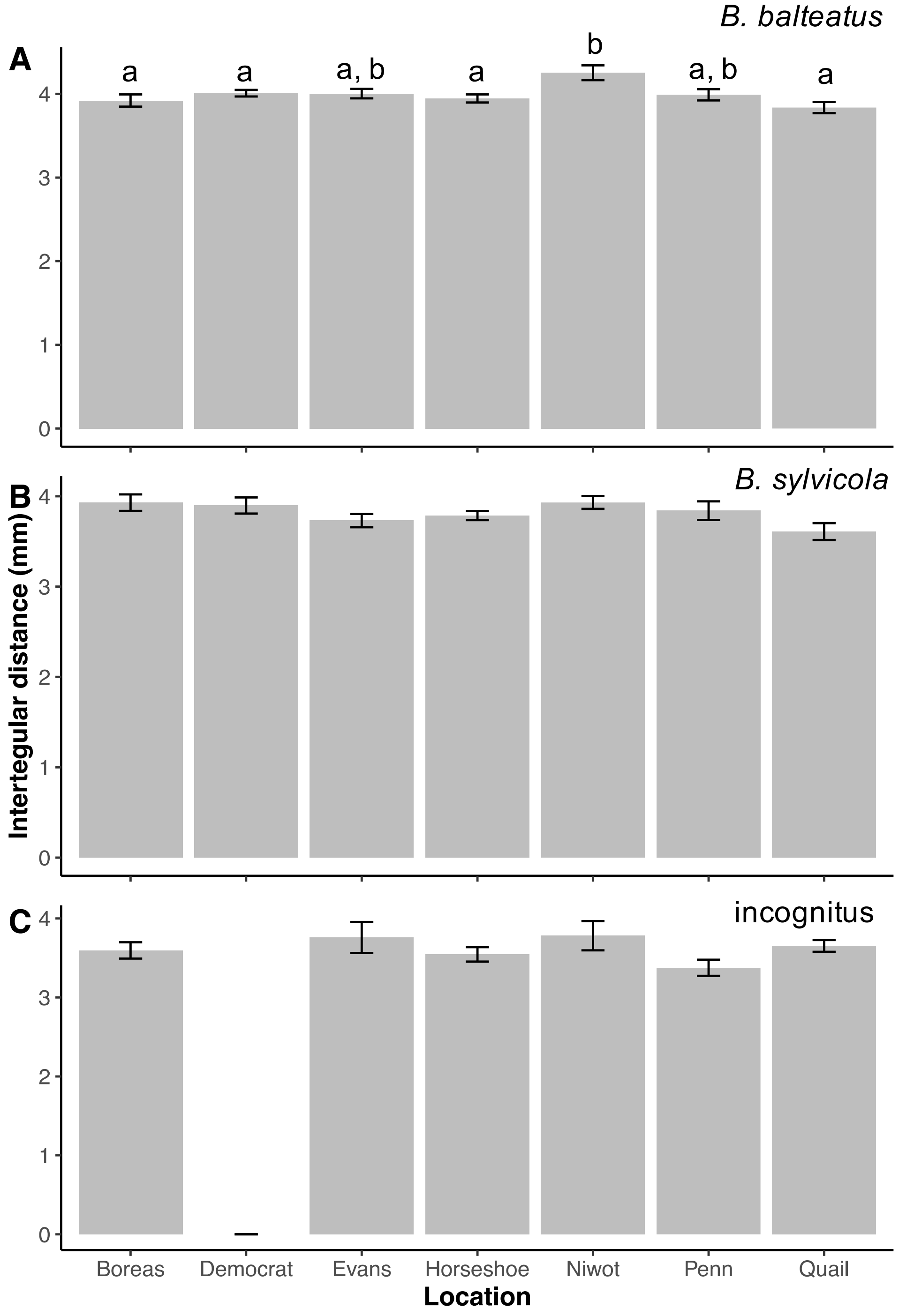


**Figure S7. Variation in intertegular distance across seven mountain locations in the Rocky Mountains for three *Bombus* species.** Average intertegular distance, a proxy for body size, for populations of (A) *Bombus balteatus*, (B) *B. sylvicola*, (C) incognitus. In each case, letters indicate significant differences between mountains based on Tukey's post hoc test, where locations not sharing letters are significantly different at $p < 0.05$. Error bars show mean ± standard error. Incognitus samples were not collected on Mount Democrat.

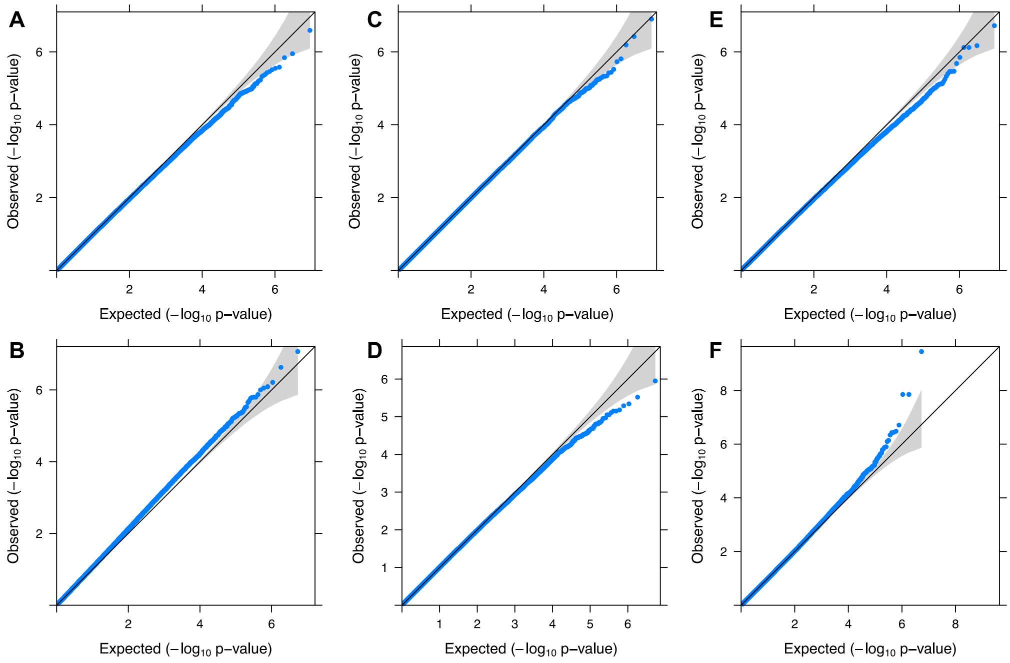


**Figure S8. Association test QQ-plots of p-values.** Observed -$\log_{10}$ p-values generated from association tests using mixed linear models between allele frequencies and phenotypes are plotted against expected values. Black diagonal represents 1:1 concordance, grey shading represents 95% confidence intervals. Phenotypes tested are (A,B) glossa length for *B. balteatus* and *B. sylvicola* respectively, (C,D) the residuals from a linear model of glossa length ~ intertegular distance to control for body size for *B. balteatus* and *B. sylvicola* respectively, and (E,F) intertegular distance for *B. balteatus* and *B. sylvicola* respectively.
